# Comparative immunogenicity of bacterially expressed soluble trimers and nanoparticle displayed influenza hemagglutinin stem immunogens

**DOI:** 10.1101/2021.03.14.435294

**Authors:** Uddipan Kar, Sara Khaleeq, Priyanka Garg, Madhuraj Bhat, Poorvi Reddy, Venkada Subramanian Vignesh, Aditya Upadhyaya, Mili Das, Ghadiyaram Chakshusmathi, Suman Pandey, Somnath Dutta, Raghavan Varadarajan

## Abstract

Current influenza vaccines need to be updated annually due to mutations in the globular head of the viral surface protein, hemagglutinin (HA). To address this, vaccine candidates have been designed based on the relatively conserved HA stem domain and have shown protective efficacy in animal models. Oligomerization of the antigens either by fusion to oligomerization motifs or display on self-assembling nanoparticle scaffolds, can induce more potent immune responses compared to the corresponding monomeric antigen due to multivalent engagement of B-cells. Since nanoparticle display can increase manufacturing complexity, and often involves one or more mammalian cell expressed components, it is important to characterize and compare various display and oligomerization scaffolds. Using a structure guided approach, we successfully displayed multiple copies of a previously designed soluble, trimeric influenza stem domain immunogen, pH1HA10, on the ferritin like protein, MsDps2 (12 copies), Ferritin (24 copies) and Encapsulin (180 copies). All proteins were expressed in *Escherichia coli*. The nanoparticle fusion immunogens were found to be well folded and bound to the influenza stem directed broadly neutralizing antibodies with high affinity. An 8.5 Å Cryo-EM map of Msdps2-pH1HA10 confirmed the successful design of the nanoparticle fusion immunogen. Mice immunization studies with the soluble trimeric stem and nanoparticle fusion constructs revealed that all of them were immunogenic, and protected mice against homologous (A/Belgium/145-MA/2009) and heterologous (A/Puerto Rico/8/1934) challenge with 10MLD_50_ mouse adapted virus. Although nanoparticle display conferred a small but statistically significant improvement in protection relative to the soluble trimer in a homologous challenge, heterologous protection was similar in both nanoparticle-stem immunized and trimeric stem immunized groups. Such rapidly producible, bacterially expressed antigens and nanoparticle scaffolds are useful modalities to tackle future influenza pandemics.

## Introduction

Influenza is a highly transmissible airborne human pathogen that can infect 5-10% of the worldwide population annually (1). Based on the surface glycoproteins Hemagglutinin and Neuraminidase, the virus can be subdivided into 18 HA subtypes and 9 NA subtypes (2,3) of which H1N1, H3N2 and an influenza B type are prevalent amongst the human population (4). Recent reports of sporadic outbreaks of H5N1 and H7N9 strains with pandemic potential in humans are a matter of concern (5)

Vaccination is considered to be the most potent strategy to check viral spread. However, the high mutational rate of viral surface glycoproteins necessitates a constant annual update of the currently prevailing trivalent as well as quadrivalent vaccines (6). Hence, several strategies have been adopted to design ‘universal influenza vaccine candidates’ that can elicit cellular as well as humoral responses superior to those triggered by a natural infection, thus providing long-lasting and cross-strain protection (7,8).

The viral surface glycoprotein Hemagglutinin (HA) is the most critical component of existing influenza vaccines (9) HA is expressed as a trimeric HA0 precursor, which undergoes cleavage to form HA1 and HA2 to convert it into a fusion active form (10). The globular HA head domain consisting solely of HA1, mediates binding to host cell sialic acid receptors while the stem domain mediates virus-host membrane fusion. Although initial studies indicated that the globular head domain is immunodominant and most neutralizing antibodies are directed towards the receptor binding site in the head domain (11), sequence analysis data revealed that the stem domain is much more conserved than the head domain (12,13). Recently, several stem-specific neutralizing monoclonal antibodies have been isolated that showed cross-neutralization against diverse influenza strains (6,14–21). This led to the idea that targeting the conserved stem domain of HA in the presence of the immunodominant HA head domain may pave the way for ‘universal influenza vaccine design’.

Application of nanotechnology has revolutionized vaccine design strategies in the past few decades including the identification of some self-assembling proteins that form nanoparticles well suited for antigen display and immune stimulation (22). The major objective was to successfully display several copies of the same immunogen on a nanoparticle platform simultaneously leading to multivalent interactions between the nanoparticle immunogen and host cell B cell receptor due to avidity effects. This should lead to enhancement in immune responses, especially B cell and T cell responses as opposed to weaker effects of monovalent interaction afforded by a single recombinant antigen. The Nabel group pioneered the design strategy by displaying repetitive arrays of trimeric HA immunogens on the three-fold symmetry axis of Ferritin that elicited antibodies with increased breadth relative to recombinant soluble HA. The antibodies generated, neutralized diverse related seasonal influenza strains but they failed to neutralize unmatched, divergent influenza strains (23). Subsequently, several groups have utilized structure-based protein design to display different immunogens on the Ferritin nanoparticle platform (24–26).

Along with Ferritin, numerous other naturally occurring proteins have the property of self-assembly that can be utilized as a multimerization platform for antigen display. Examples of such other multimeric display platforms include MsDps2 (PDB-2Z90), a dodecameric DNA binding protein from stationary-phase cells of *Mycobacterium smegmatis* that has four three-fold axes of symmetry (27) and Encapsulin (PDB-4PT2), a 180-mer iron storing protein of *Myxococcus xanthus* (28). Current nanoparticle designs typically have either or both of the nanoparticle scaffold and the displayed antigen expressed in mammalian cells. This limits rapid scale up and deployment, especially in low resource settings.

Previously, we reported the design of a headless stem domain immunogen H1HA10-Foldon based on H1N1 A/Puerto Rico/8/34 that was bacterially expressed and conferred complete protection against homologous challenge in a mice model (29). With this background, we attempted to display pH1HA10 derived from H1N1 (A/California/04/2009) (29) on MsDps2 (PDB-2Z90), Encapsulin (PDB-4PT2) and Ferritin (PDB-3BVE) protein nanoparticles and compare with corresponding soluble trimeric antigen, pH1HA10-Foldon.

The designed nanoparticle immunogens were found to be properly folded and exhibited strong binding to the stem-directed antibodies thus validating the design strategy. In addition, we obtained a cryo-EM map of MsDps2 displaying the influenza stem immunogen (pH1HA10 on MsDps2 protein nanoparticles) at 8.5 Å where the MsDps2 core is resolved at 3.45 Å. In mice, following intramuscular immunization with oil-in-water adjuvanted formulations challenge studies demonstrated that both Msdps2-pH1HA10 and Encapsulin-pH1HA10 could confer complete protection against both homologous (Bel 09) as well as heterologous (PR8) viral challenge. In comparison, the soluble pH1HA10-foldon trimer conferred 80% and 100% protection against homologous and heterologous challenge, respectively.

## Materials and Methods

### Design of MsDps2-pH1HA10, Encapsulin-pH1HA10 and pH1HA10-Ferritin constructs

The previously reported stem immunogen pH1HA10 (29) was fused at the C termini of MsDps2 (residues 1-161) (source: *Mycobacterium smegmatis*) (accession number A0QXB7) and Encapsulin (residues 1-287) (source: *Myxococcus xanthus*) (accession number Q1D6H4) and at the N terminus of Ferritin (residues 1-163) (source: *Helicobacter pylori*) (accession number Q9ZLI1). The immunogen was connected to the scaffold using linkers of 9 residues (GSAGSAGAS), 15 residues (GSAGSAGSAGSAGAS) and 5 residues (QASGG) respectively. MsDps2-pH1HA10 and pH1HA10-Ferritin were in frame with an N terminus 6x Histidine tag followed by thrombin cleavage site for Immobilized Metal Affinity Chromatography (IMAC) Ni-NTA resin purification. Encapsulin-pH1HA10 contained a C-terminal 6x Histidine tag. The nanoparticle-stem fusion constructs were codon optimized for expression in bacteria and synthesized by GenScript (New Jersey, USA). The constructs were cloned in the expression vector pET-28a (+) (Novagen) in between the Nde1 and Hind III sites. The synthesized plasmids were purified, and restriction digestion (both single and double digest) was done to confirm the insert size. pH1HA10-Foldon was rationally designed based on the crystal structure of H1N1 A/California/04/2009 (PDB-3LZG). The CR6261 epitope was mapped onto Cal09 HA and fragments covering 80% of the epitope with stable breakpoints were selected and joined through linkers of suitable length. Hydrophobic residues generated during the process, due to loss of interactions with the rest of HA, were resurfaced. Additional mutations were incorporated to retain the HA stem in a neutral pH conformation (30). Trimerization was accomplished by fusion of a T4 bacteriophage derived ‘Foldon’ domain at the C terminus of pH1HA10, as reported previously (29).

### Expression and purification of the stem- nanoparticle fusion proteins

The nanoparticle-stem fusion constructs were purified from *E. coli* BL21 (DE3) cells following T7 promoter induction. Briefly, a single sequence confirmed colony from the transformed plate was first inoculated in 5 mL of primary culture and grown overnight at 37 °C. This was followed by secondary inoculation of 1% of primary culture in 500 mL Terrific Broth media (Himedia Laboratories Pvt. Ltd.) which was then grown at 37 °C till an OD_600_ of 0.6-0.8 was reached. The culture was then induced with 1 mM IPTG. After inducing the cells for 12-16 hours at 20 °C, the cells were harvested through centrifugation at 4000 rpm, 4 °C for 15 minutes, followed by sonication in 1x PBS (pH 8). A second round of centrifugation at 12,000 rpm, 4 °C for 20 minutes, was then carried out to separate the soluble fraction from the insoluble fraction of the cell lysate. The nanoparticle-stem fusion constructs were purified from the inclusion bodies. The cell pellet was solubilized in 1x PBS buffer (pH 8), supplemented with 150 mM NaCl and 6 M Guanidine Hydrochloride. The soluble fraction was then allowed to bind to Ni-NTA resin (GE Healthcare Life Sciences, Amersham) at 25 °C and eluted in a gradient of imidazole (100 mM, 200 mM, 300 mM, and 500 mM) in 1x PBS buffer (pH 8), containing 150 mM NaCl and 6 M Guanidine Hydrochloride. Pooled, eluted fractions were refolded *in vitro* by gradual removal of Guanidine Hydrochloride through 3 rounds of dialysis against 2 L each of the refolding buffers as follows-Round 1: 1 M Guanidine Hydrochloride supplemented with 400 mM Arginine Hydrochloride, 1x PBS (pH 8), round 2-1x PBS (pH 8) containing 100 mM Arginine Hydrochloride and finally, round 3-1x PBS (pH 7.4) with 10% glycerol for storage.

### SDS- PAGE and Circular Dichroism

Protein purity was assessed through denaturating and reducing SDS-PAGE. Samples were boiled in Dithiothreitol containing SDS sample loading buffer. Thermal unfolding of Encapsulin-pH1HA10 was followed by recording the CD spectra on a JASCO J-715 spectropolarimeter. 8 μM of the protein in 1x PBS (pH 7.4) was used. Spectral measurement was done at 222 nm in a temperature range of 10 °C to 90 °C, with a 10 °C/min rise. Data acquisition was done at 25 °C with a 0.1 cm path length quartz cuvette, response time of 16 seconds, and a spectral bandwidth of 2 nm. Normalized mdeg was plotted as a function of temperature.

### Size Exclusion Chromatography (SEC), and Size Exclusion Chromatography –Multi Angle Light Scattering (SEC-MALS)

To determine the oligomeric status of the purified fusion proteins, size exclusion chromatography (SEC) was performed in non-denaturing conditions. 500 μL of the proteins ranging from 2 mg/mL – 4 mg/mL concentration were loaded on an analytical Superose 6 Increase 10/300 column (GE Healthcare) and eluted in 1x PBS (pH 7.4) at 0.4 mL/min flow rate on an Äkta pure chromatography system. A low molecular weight gel filtration calibration kit (product code-28403841) (GE Healthcare) was used for calibrating the column. For SEC-MALS (SEC-Multiangle light scattering), proteins were separated on a Superose 6 Increase 10/300 column (GE Healthcare) in 1x PBS (pH 7.4) at a flow rate of 0.4 mL/min. SEC resolved peaks were subjected to an in-line MALS detector (mini-DAWN TREOS, Wyatt Technology corp.) and a refractive index monitor (WATERS corp.) for molecular weight estimation. The data was analyzed through ASTRA 6.0 software (Wyatt Technology), as described previously (31).

### Binding affinity studies by Surface Plasmon Resonance (SPR)

The binding affinity of all fusion constructs to the stem-directed broadly neutralizing antibodies (bnAbs) CR6261 and CR9114 was measured by SPR studies performed on a Biacore 3000 optical biosensor (Biacore, Uppsala, Sweden) at 20 °C. Ligand immobilization was done through amine coupling to an NHS-EDC activated CM5 chip (GE HealthCare, Uppsala, Sweden). CR6261 or CR9114 were immobilized at 1000 RUs in 10 mM Sodium Acetate buffer (pH 4.5) at a flow rate of 30 μL/min. One channel was left blank to act as a reference channel. The analyte, nanoparticle fusion constructs were passed in a concentration series over ligand immobilized sensor channels in PBS (pH 7.4) with 0.05% P20 surfactant. A flow rate of 30 μL/min was maintained throughout the binding interactions. The sensor surface was regenerated after use for successive experiments by passing 4 M magnesium chloride over the chip surface. The final trace was corrected for non-specific binding through subtraction of the interaction signal from the ligand-free blank channel. The data was processed and global fitting with a simple 1:1 Langmuir interaction model was done using BIA-evaluation 3.0.1 software to obtain the kinetic parameters.

### SPR Binding of proteins to stem directed broadly neutralizing antibody CR9114, following thermal stress

The proteins in 1x PBS (pH 7.4) were exposed to transient thermal stress by incubating at a range of temperatures from 20 °C to 80 °C in a thermal cycler (Eppendorf) for 1 hour. Subsequently, the proteins were returned to room temperature and binding to CR9114 at 100 nM concentration of the proteins was monitored in a Biacore 3000 SPR instrument (Biacore, Uppsala, Sweden) as mentioned in the previous section.

### Binding affinity studies by Microscale Thermophoresis (MST)

MST was performed as an alternative approach to measure affinities (32,33). In this assay, bnAb CR6261was labelled using a Monolith™ Protein Labelling Kit RED-NHS (Catalogue Number-MO-LO11) (Nano Temper Technologies) in the supplied labelling buffer as per instructions in the manufacturer’s manual. Following that, the protein was eluted into an assay buffer for MST (specifically buffer A including 0.05% Tween 20). Non-fluorescent immunogens with concentrations ranging between 5 μM and 1 pM were titrated in a two-fold dilution series and mixed with a fixed amount of labelled Ab, here CR6261 (100 nM). Next, MST Premium Coated (NanoTemper Technologies) Monolith™ NT.115 capillaries were loaded with the samples and measurement was done with the help of Monolith NT.115 and MO. Control software at room temperature at different power intensities (LED/excitation power setting 80%, MST power setting medium and high). Finally, MO.Affinity Analysis software (version 2.2.5, NanoTemper Technologies) was used to analyze the data (34).

### Thermal Melt studies using nanoDSF

Equilibrium thermal unfolding of MsDps2-pH1HA10 and pH1HA10-Ferritin was carried out using nanoDSF (Prometheus NT.48). 4 to 5.5 μM of the proteins in 1x PBS (pH 7.4) were incubated in the temperature range of 20 °C to 80 °C at 50% LED power, as previously described (35).

### Negative stain single particle EM (Electron Microscopy)

All the negative stain EM samples were prepared using staining methods described in previous studies (36,37). Initially, glow discharge of the copper grids was carried out for 40-60 seconds at 20 mA current using GloQube glow discharge system (Quorum technologies) for uniform settling of the particles on the grid. Following this, 3.5 μl of nanoparticle (0.4 mg/mL) in 20 mM Tris was adsorbed on the grids for 30-60 secs. Excess buffer was removed by blotting with a filter paper and negative stain was performed using 1% uranyl acetate. Negative stain imaging experiment was carried out at room temperature using a FEI Tecnai 12 BioTwin transmission electron microscope equipped with a LaB6 (lanthanum hexaboride crystal) filament at a voltage of 120 Kv. Datasets were collected using a side-mounted Olympus VELITA (2Kx2K) CCD camera at a calibrated magnification of 75000 x and a defocus value of −1.3 μm. The final images were recorded at a pixel size of 2.54 Å on the specimen level. Following this, properly sized particles were manually chosen and extracted using e2boxer.py and e2projectmanager.py EMAN 2.1 software (38). Finally, a 2-D reference free classification of particles was carried out using e2refine2d.py.

### Cryo-EM sample preparation and data acquisition

Quantifoil R2/1 or R1.2/1.3 300 mesh holey carbon grids were glow discharged for 2 minutes at 20 mA. The freshly prepared 3 μL protein samples at a concentration of 0.5 mg/ml were added to the carbon grids and incubated for 10 seconds before plunging into liquid ethane using FEI Vitrobot Mark IV plunger at 20 °C with 100% humidity. The grids were blotted for 3-5 seconds at zero blot force. Cryo-EM data acquisition of the frozen sample was carried out using a Thermo Scientific™ Talos Arctica transmission electron microscope at 200 kV at a nominal magnification of 42,200 x, equipped with a K2 Summit Direct Electron Detector. LatitudeS automatic data collection software (Gatan Inc) was utilized to collect the final 746 digital micrographs at a pixel size of 1.2 Å with a total electron dose of about 40 e-/Å^2^ at the defocus range of −1.25 and −3.5 μm, under a calibrated dose of about 2 e-/Å^2^ per frame. Movies were recorded for 8 secs with 20 frames. All the collected micrographs were then motion corrected using “dosefgpu_driftcorr” using MotionCor2 software (39).

### Cryo-EM data processing and 3-D reconstruction

Relion 3.0 was used to process the whole dataset (40,41). The beam-induced motion correction of the individual movies was performed using MotionCor2 software. Following this, cisTEM software package (38) was used to determine the quality of the motion corrected micrographs. The contrast transfer function (CTF) correction was carried out using CTFFIND4 (42). A total of 4,29,709 particles were automatically selected using Relion auto-picking. The full data set was binned by 2X (2.4 Å) for reference-free 2D classification with a box size of 256 pixels and particle diameter of 220 Å. After three rounds of 2D classification, about 3,14,457 particle projections were selected for further 3D classification. Additionally, a small dataset, around 10,000 particle projections were selected for calculation of the de novo initial model. This initial model was low pass filtered to 40 Å and 3,14,457 particle projections were selected for 3D classification with T symmetry imposed, and the whole data set was split into 6 classes. For this condition, a soft mask (220 Å) was implemented to mask out the immunogen densities. Only non-bin data (1.2 Å) was utilized to calculate the final 3D structure. The best classes with stable connection between MsDps2 core and pH1HA10 extension were merged for further refinement. The 3-D reconstruction map was further subjected to focus-based refinement to separately focus on only the MsDps2 nanoparticle core (using a tight mask of 160 Å). These particles were subjected to another round of refinement for CTF and beam tilt in RELION 3.0. Subsequent movie-refinement was followed by particle polishing for further refinement. The two unfiltered half-maps were used to calculate the resolution, and the resolution of the 3D model of MsDps2 fusion was estimated at 0.143 of Fourier shell correlation (FSC) (43).

### Docking of the crystal structure into the Cryo-EM map of MsDps2-pH1HA10 and subsequent refinement

Initially, UCSF Chimera (44) was used to visualize the Cryo-EM map as well as the available crystal structure of MsDps2 (PDB-2Z90) (27). pH1HA10 stem domain immunogen was modelled on the crystal structure of HA from the H1N1 A/California/04/2009 strain (PDB-3LZG) (45) as described previously (29). Briefly, the antibody footprint of CR6261 was mapped onto the structure of the full length H1N1 A/California/04/2009. HA stems fragments covering ~80% of the CR6261 epitope with stable breakpoints and optimal termini distance were identified and connected by linkers of suitable length. Newly generated hydrophobic surface exposed residues, due to loss of interaction with the rest of the HA, were mutated. Low-pH conformation destabilizing mutations were also incorporated to retain the HA stem in the neutral-pH conformation. The modelled structure of pH1HA10 stem domain immunogen as well as the coordinates of MsDps2protein crystal structure (PDB-2Z90) were manually fit into the cryo-EM map of MsDps2-pH1HA10 using the “fit in Map” command of UCSF Chimera. The final docking was performed using the automatic fitting option of the UCSF Chimera visualization package.

### Immunization studies in Mice

All the immunization experiments were carried out by Mynvax Pvt. Ltd, India (Study site: Central Animal Facility (CAF), IISc). BALB/c mice (n= 5/group, female, 6-7 weeks old, 16-19 g) were vaccinated with freshly adjuvanted (Sepivac SWE (Cat. No. 80748J, Batch No. 200915012131, SEPPIC SA, France) protein with a dose of 20 μg/ animal per dose of nanoparticle-stem fusion proteins (1:1 v/v antigen: adjuvant, 20 μg in 50 μL 1x PBS (pH 7.4) and 50 μL Adjuvant) via intramuscular route in a prime and boost regime on Day 0 and Day 21, respectively. Sera was collected by bleeding the animals at Day −1 (pre-bleed), Day 14 (post prime) and Day 35 (post boost) through retro-orbital puncture. The immunization study performed was approved by Institutional Animal Ethics Committee (IAEC) No. CAF/ETHICS/848/2021. **Viral Challenge**

For viral challenge, immunized mice were housed in individually ventilated cages; ambient room temperature 22±2 °C, relative humidity 60±10%, 12 h light and dark cycle was maintained, for viral challenge. 2 weeks post the boost immunization, animals were challenged with either 10423 pfu (10MLD_50_) of antigen matched strain (H1N1 A/Belgium/145-MA/2009) (homologous challenge, n= 5/group, 10423 pfu) (kindly provided by Dr. Xavier Saelens, VIB-UGent Center for Medical Biotechnology, Belgium) or 270 pfu (10MLD_50_) of antigen mismatched strain (H1N1 A/Puerto Rico/8/1934) (heterologous challenge, n=5/group, 270 pfu) (kindly provided by DR. Shashank Tripathi, Center for Infectious Disease Research, IISc, India) intranasally after anesthetizing with ketamine and xyylazine cocktail (2 mg ket/0.2 mg Xyl/ 20 g mice) intraperitoneally. Body weight and temperature were recorded upto two weeks post challenge. Animals with body weight reduction equal to or lower than 25% were euthanized humanely. Clinical signs were measured as: no abnormality detected (0 points), mild piloerection (1 point), piloerection (2 points), shivering or severe shivering (1 point), emaciation (1 point) and hunching (1 point), upto two weeks post challenge.

### ELISA studies

ELISA was carried out with the post boost sera to determine the titer of antibodies elicited by the stem-nanoparticle immunogens. Desired antigens (4 μg/ml in 1x PBS (pH 7.4), 50 μL/well) were coated on 96-well Nunc plates (Thermo Fisher Scientific, Rochester, NY) at 4 °C overnight under constant shaking (300 rpm). After washing with PBST (PBS having 0.05%Tween-20, 200 μL/well), plates were blocked using blocking solution (3% skim milk in PBST) and incubated at 25 °C for 1 h, 300 rpm. Post blocking, fourfold serial dilutions of sera starting at 1:100 dilution were added (50 μL/well) and incubated at 25 °C for 1 h, 300 rpm. The plates were again washed with PBST (200 μL/well). Next, ALP-conjugated goat anti-mouse IgG secondary antibody (Sigma-Aldrich) in PBST was added at a predetermined dilution (1:5000 in blocking buffer) (50 μL/well) and incubated at 25 °C for 1 h, 300 rpm. Following a final round of wash with PBST (200 μL/well), pNPP substrate (pNPP, Sigma-Aldrich) was added (50 μL/well) and incubated at 25 °C for 30 min, 300 rpm. The choromogenic signal was read at 405 nm. End point titer was calculated as the highest serum dilution giving signal above cutoff (0.2 O.D. at 405 nm).

### Microneutralization assay

Viruses were grown in Madin-Darby Canine Kidney (MDCK) cells in the presence of TPCK-treated trypsin (4 μg/mL) and stored at −70 °C. Immune mice sera samples were heat-inactivated and treated with receptor destroying enzyme (RDE, SIGMA-ALDRICH) before use. Immunized sera samples were two-fold serially diluted and incubated for 1 h at 37 °C in 5% CO_2_ with 50 TCID_50_ viruses. Serum-virus mixture was then transferred to 96 well plates, and 1.5 × 10^5^ MDCK-London cells/mL were added to each well. Following which plates were incubated for 48 h at 37 °C in 5% CO_2_, and cytopathic effects were observed. The neutralization titer in the assay is the highest serum dilution at which no cytopathic effect was observed, as described previously (46). Two-tailed Mann Whitney *t*-test was performed for pairwise MN titer comparisons.

### Competition assay with CR6261 using SPR

Competition between the antigen elicited sera and bnAb CR6261 was checked through SPR on a Biacore 3000 optical biosensor (Biacore, Uppsala, Sweden) at 20 °C. Competition was studied using pH1HA10-Foldon immobilized on an NHS-EDC activated CM5 chip (GE HealthCare, Uppsala, Sweden) through amine coupling until 1500 RUs were achieved and subsequently probing with two analytes without an intermittent dissociation step. First, 3-fold dilutions of the pooled sera of each antigen were passed for 100 secs (flow rate: 30 μL/min) starting at their pH1HA10-Iz binding GMT, which was evaluated in the previous section through ELISA. Subsequently, 5 μM of CR6261 (saturating concentration of the antibody which was evaluated separately by titrating varying concentrations of the antibody against the immobilized, trimeric stem) was flowed at 30 μL/min and association was observed for 100 secs. Finally, dissociation was followed for 200 secs. Competition of the sera antibodies with CR6261 for binding to overlapping epitopes on the trimeric stem was estimated by comparing the change in RUs observed for CR6261 binding to pH1HA10-Foldon in the absence or presence of sera. To rule out non-specific competition by sera components, PBS immunized mice sera was used as control. CR6261 binding with immobilized stem was used as binding control in the absence of sera. All proteins were diluted in 1x PBS (pH 7.4). The percent competition was calculated using the following equation (equation 1):

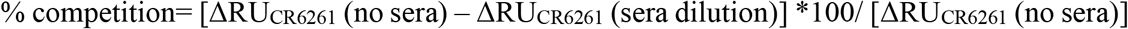

where, ΔRU_CR6261_ (no sera) is the change in RUs (from 0 secs to 100 secs) of CR6261 binding to pH1HA10-Foldon in the absence of sera, and ΔRU_CR6261_ (sera dilution) is the change in RUs (from 0 sec to 100 sec) of CR6261 binding to pH1HA10-Foldon in the presence of various dilutions of sera. IC_50_ value was calculated as the dilution at which 50% competition of the sera with CR6261 was observed for binding to pH1HA10-Foldon.

### Statistical Analysis

Statistical analysis for ELISA binding titer assays was done using a two-tailed Mann-Whitney t test inbuilt in the GraphPad Prism software 9.0.0. The p values are as follows-ns indicates not significant, * indicates p<0.05, ** indicates p<0.01, **** indicates p< 0.0001. Statistical significance of survivability between vaccinated and unvaccinated groups was compared using two-way ANOVA test with Bonferroni correction. (ns= not significant, p < 0.01 (*), p < 0.001 (**), p < 0.0001 (****)). Multiple T test with Bonferroni Dunn correction was used to compare body weight reduction between groups (p < 0.01 (*), p < 0.001 (**), p < 0.0001 (****)). Microneutralization assay between different groups were compared using two-tailed Mann Whitney t test (ns indicates not significant, * indicates p < 0.05, ** indicates p < 0.01, and **** indicates p < 0.0001).

## Results

### Design of nanoparticle-HA stem fusion immunogen

Multimerization of antigens to mimic the highly repetitive and structurally ordered arrangement of antigenic proteins on the viral surface can induce robust titers of binding and neutralizing antibodies (47–51). Multi-component nanoparticle platforms activate the adaptive immune system by stimulating B and T cell responses (52,53). Previously, the Nabel group successfully displayed trimeric influenza immunogens on Ferritin nanoparticles which induced neutralizing antibodies (23). Using a similar approach, we attempted to utilize an iterative structure-guided design strategy to display the previously described HA stem immunogen, pH1HA10-Foldon, from H1N1 A/California/04/2009 (29) on self-assembling protein nanoparticle platforms. We fused the pH1HA10 stem immunogen either to the C termini of MsDps2 (*PDB id*. 2Z90) or Encapsulin (*PDB id*. 4PT2) or to the N terminus of Ferritin (PDB id. 3BVE) (Figure 1A, Table 1). Since geometrical constraints in the 12-meric MsDps2, and 24-meric Ferritin enforce three-fold symmetry, the trimerization domain, Foldon, was removed. We hypothesized that pH1HA10 could be displayed as a trimer on both MsDps2 and Ferritin. In contrast, Encapsulin, which has a six-fold axis of symmetry, could display 180 molecules of the monomeric stem immunogen. Previous studies have shown that monomeric stem immunogens also elicit protective antibodies (29). Using these various display strategies would also enable comparison between the choice of nanoparticles as scaffolds for natively oligomerized antigens. Relative surface area (RSA) calculations indicated that fusion between the solvent exposed C termini of MsDps2 and Encapsulin and N terminus of Ferritin to the available N/C termini of pH1HA10 was feasible. The coaxially aligned planes containing the fusion termini of the stem and the nanoparticle were separated to prevent steric clashes. The distance between the N terminus of pH1HA10 and the C terminus of MsDps2 or Encapsulin was 21 Å and 34 Å respectively, while the distance between the C terminus of pH1HA10 and N terminus of Ferritin was 12 Å (Table 1) (Figures 1A-B). A representative design scheme for MsDps2-pH1HA10 is shown in Figure 1B. Consequently, MsDps2, Encapsulin and Ferritin were fused to the stem through 9 residue (GSAGSAGAS), 15 residue (GSAGSAGSAGAS) and 5 residue (QASGG) long flexible linkers, respectively. In summary, the nanoparticle-stem immunogen had the HA stem fused to either the N or C terminus of nanoparticles through a linker, with a cleavable N or C terminal 6x Histidine tag for Immobilized Metal Affinity Chromatography (IMAC) Ni-NTA purification (Figure 1C).

**Figure 1:**
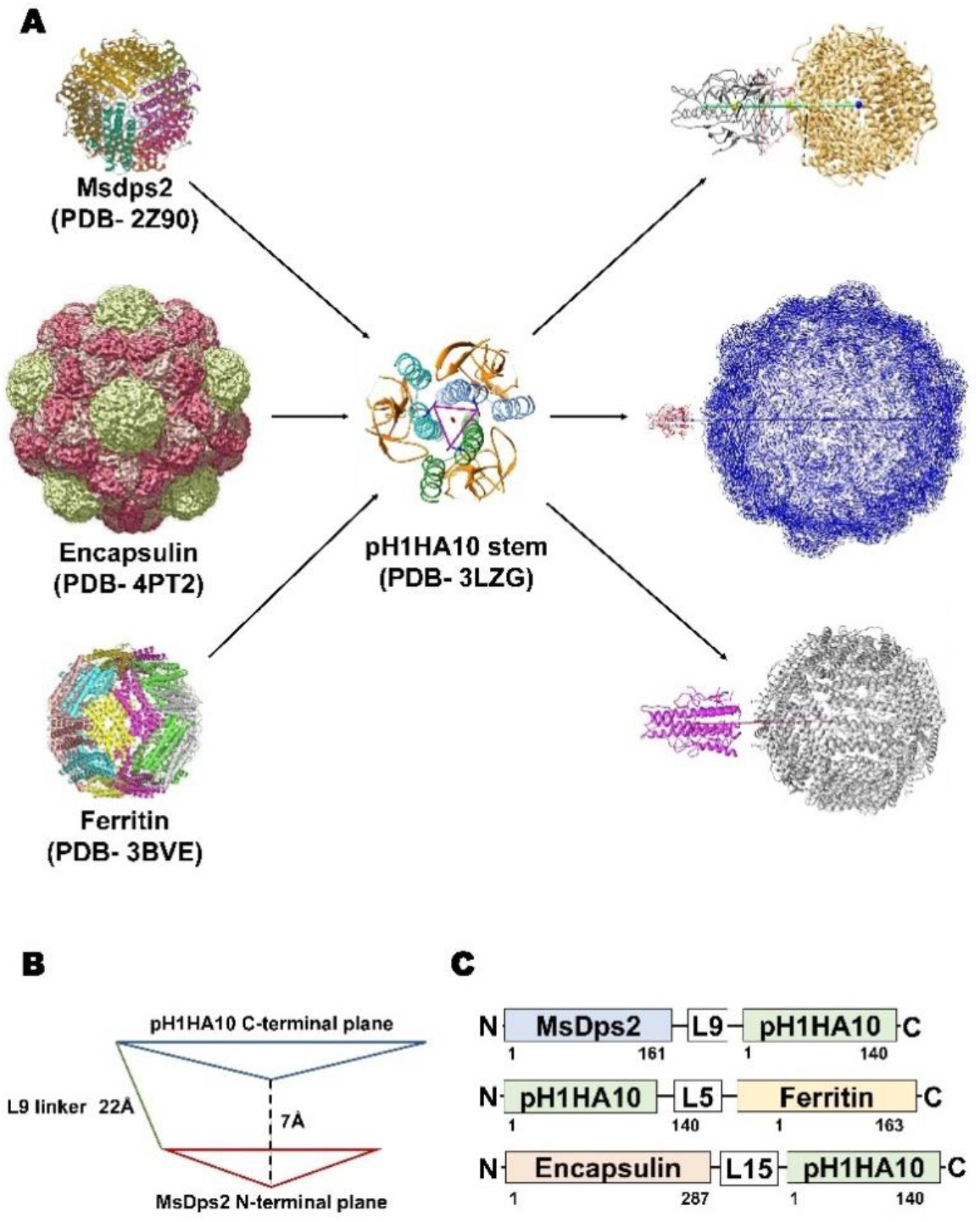
Design of pH1HA10-nanoparticle fusion constructs. **(A)** Representative diagram showing stem immunogen, pH1HA10, displayed on the various sized nanoparticle platforms, Msdps2 (PDB id. 2Z90) Encapsulin (PDB id. 4PT2) and Ferritin (PDB id. 3BVE). **(B)** Schematics of construct design for MsDps2-pH1HA10. The coaxially aligned planes containing the C and N terminal, respectively, of MsDps2 and pH1HA10 are separated by 7Å to avoid steric hindrance. The distance between the N terminal Asp1 of pH1HA10 and C terminal Val161 of MsDps2 is 21Å, which is spanned by a 9-residue long linker. The same design was extended to Encapsulin and Ferritin-stem fusion contructs. **(C)** MsDps2 and Encapsulin are linked to the N terminus of pH1HA10 through 9-residue and 15-residue long linkers (L9 and L15) respectively, whereas Ferritin is fused at the C terminus of pH1HA10 through a 5-residue long linker (L5).

**Table 1:**
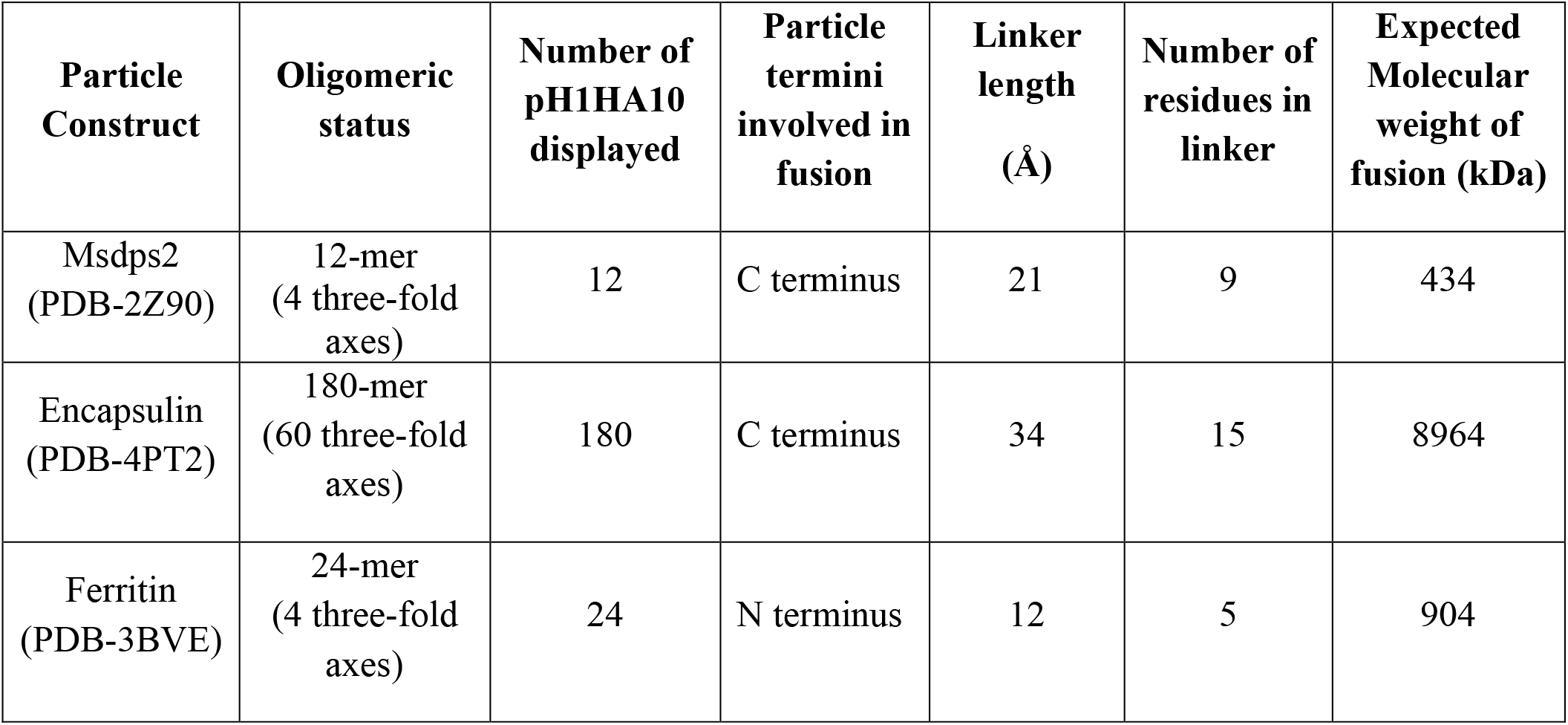
Details of the nanoparticle platforms used:

### Purification and biophysical characterization of designed pH1HA10-nanoparticles

We sought to express these nanoparticle-stem immunogens in bacteria because of the rapid scalability and cost-effective nature of bacterial systems. Consequently, all the fusion proteins were expressed in *E. coli* BL21 (DE3) and purified and refolded from inclusion bodies. Yield and purity were confirmed by SDS-PAGE analysis (Figures 2A-C).

**Figure 2:**
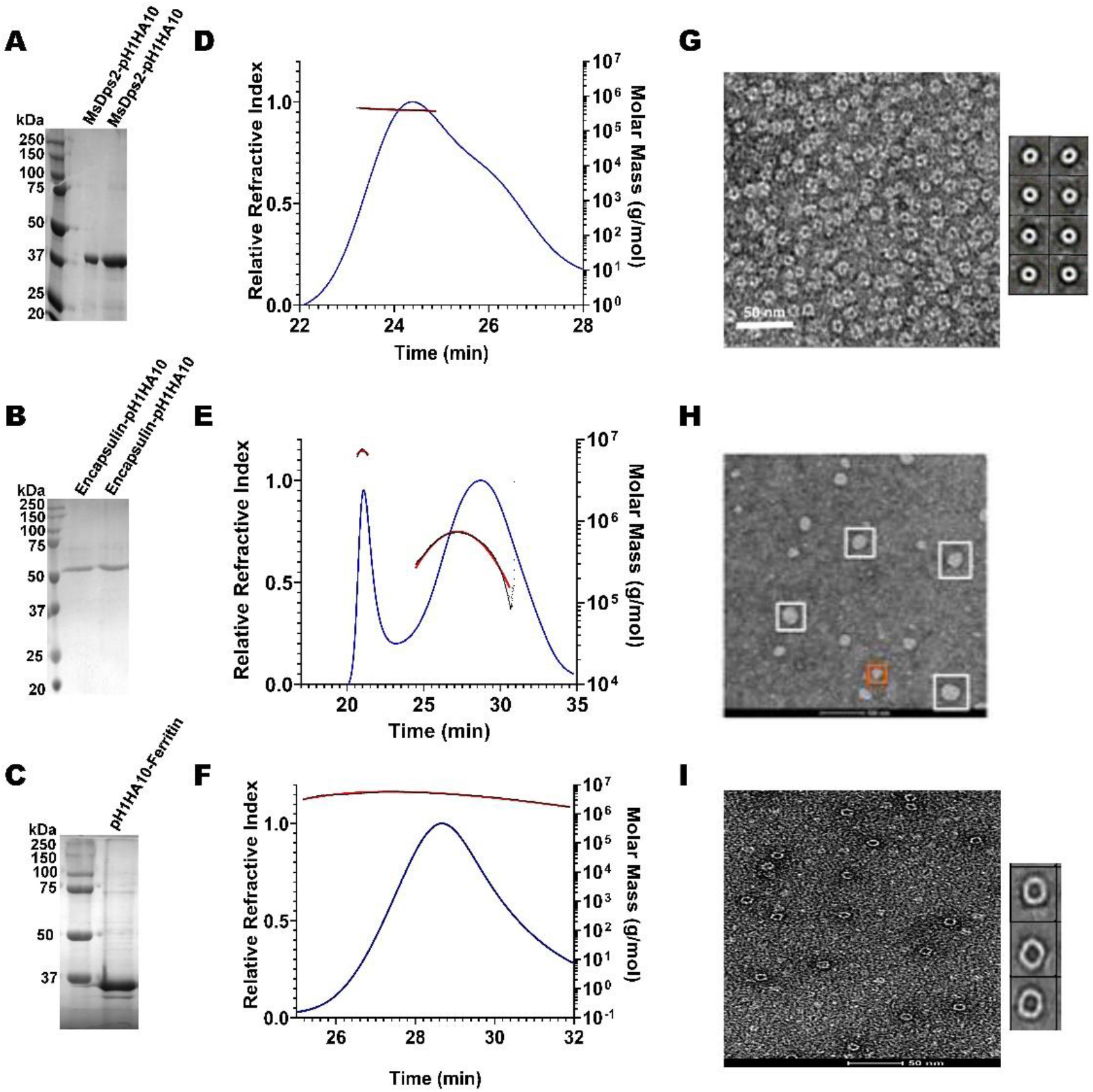
Biophysical characterization of pH1HA10 fused nanoparticles. **(A-C)** SDS-PAGE analysis of purified stem-nanoparticles- **(A)** MsDps2-pH1HA10 (MW: 36.1 kDa), **(B)** Encapsulin-pH1HA10 (MW: 49.7 kDa) and **(C)** pH1HA10-Ferritin (MW: 38.6 kDa) under reducing conditions. **(D-F)** SEC-MALS profile of **(D)** Msdps2-pH1HA10, **(E)** Encapsulin-pH1HA10, and **(F)** pH1HA10-Ferritin confirm the formation of oligomers of 12-mer, 180- mer and 24- mer, respectively. **(G-I)** Negative stain EM images of **(G)** MsDps2-pH1HA10 with left panel showing micrograph of negative stain EM images for MsDps2-pH1HA10, and the right panel showing 2D class averages obtained from the EM data. **(H)** Encapsulin-pH1HA10 forms heterogenous mixture of species in solution. **(I)** pH1HA10-Ferritin exists as homogenous oligomers as shown in negative stain image (left) and 2D class average (right).

Degree of oligomerization and conformational heterogeneity of the designed nanoparticle fusion protein was confirmed by analytical size exclusion chromatography (SEC) under native conditions as well as negative staining TEM imaging at room temperature MsDps2-pH1HA10 existed as homogenous dodecamers in solution and the molar mass was estimated from SECMALS to be 428.5 ± 2.4 kDa, whereas, Encapsulin-pH1HA10 distributed into a heterogenous mixture of oligomeric species, with a predominant 180-meric protein of molar mass estimated to be 8756 ± 40 kDa and a smaller oligomeric species with an average molar mass of 992 ± 10 kDa. pH1HA10-Ferritin also formed a homogenous population of 24-meric protein in solution with an observed molar mass of 979 ± 39 kDa (Figures 2D-F). EM images and 2D reference free class averages of MsDps2-pH1HA10 and pH1HA10_Ferritin also indicated a uniform spherical particle distribution, which closely resemble the unconjugated MsDps2 or Ferritin proteins indicating that fusion to the influenza immunogen does not alter the overall morphology of the nanoparticle core. Encapsulin-pH1HA10 formed poly-dispersed spherical particles of varying diameter (Figure 2G-I). However, the displayed pH1HA10 immunogen on the nanoparticle platforms was unnoticeable in negative staining TEM images. The flexible nature of the linker connecting the immunogen to the nanoparticle is likely averaged out during the 2D classification making the immunogen apparently invisible. The size of the immunogen is small compared to the core nanoparticle, which also allows one average out small immunogen densities from the core in 2D class averages. However, in the Cryo-EM structure of MsDps2-pH1HA10, solved by us, the stem spikes on the nanoparticle were clearly visible.

### Binding studies with stem directed broadly neutralizing antibodies (bnAbs)

Our stem immunogen mimics the pre-fusion, neutral pH conformation of full-length HA which contains conformational epitopes for a Group 1 specific stem directed antibody, CR6261. We previously showed that the monomeric stem, H1HA10, bound CR6261 with sub micromolar affinity (K_D_: 315.4 ± 14.5 nM) and a soluble trimeric version of the stem immunogen, H1HA10-Foldon, bound CR6261 with a K_D_ of 52.4 ± 1.8 nM (29). To ascertain proper folding of antigen display on nanoparticles, we examined the binding affinity of stem-nanoparticle fusions with the stem-directed conformational epitope specific antibody CR6261. MsDps2-pH1HA10 and Encapsulin-pH1HA10 bound CR6261 with 16-fold (K_D_: 3.2 ± 0.5 nM) and 105-fold (K_D_: 0.5 ± 0.01 nM) higher affinity, respectively whereas, pH1HA10-Ferritin bound without any significant dissociation, compared to the trimeric immunogen (Figures 3A-B, E). MsDps2-pH1HA10 and Encapsulin-pH1HA10 bound pan-influenza (neutralizes both group 1 and group 2) stem specific bnAb, CR9114, with KD of 4.4 ± 0.3 nM and 5.5 ±1.3 nM respectively (Figures 3C-D).

**Figure 3:**
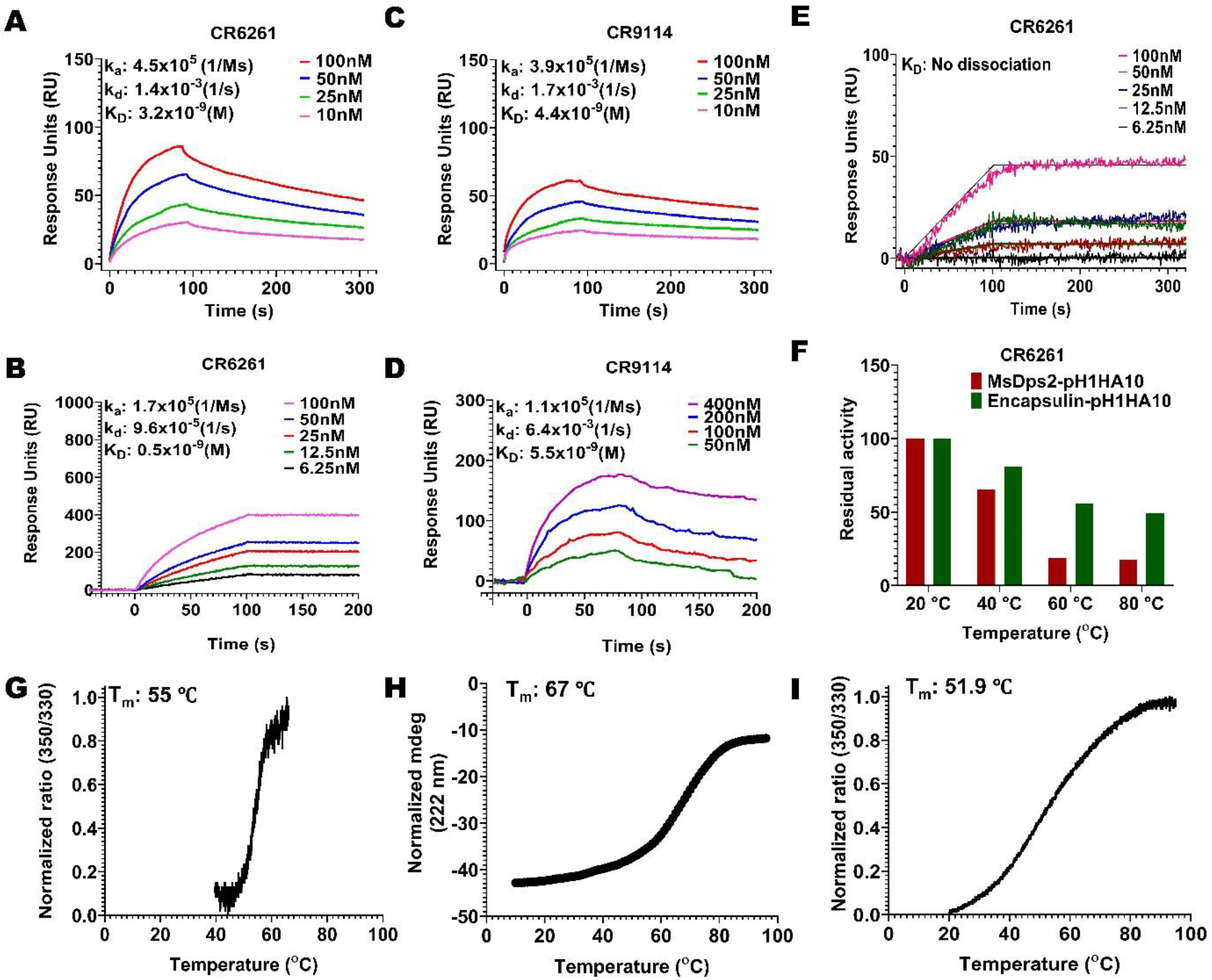
SPR studies and thermal stability of pH1HA10-nanoparticle immunogens. **(A-D)** SPR binding studies of MsDps2- pH1HA10 and Encapsulin-pH1HA10 with CR6261 **(A-B)** or CR9114 **(C-D)**. **(E**) SPR binding of pH1HA10-Ferritin with CR6261. **(F)** Thermal tolerance of MsDps2- pH1HA10 and Encapsulin- pH1HA10. Proteins were incubated at indicated temperatures for 1h and returned to RT for SPR binding with CR6261. **(G)** nanoDSF monitored thermal denaturation profile of MsDps2-pH1HA10. **(H)** CD melt profile of Encapsulin-pH1HA10. **(I)** nanoDSF monitored thermal melt of pH1HA10-Ferritin.

For further validation, the binding of the nanoparticle fusion constructs to CR6261 antibody was characterized using Microscale thermophoresis (MST) and K_D_’s obtained were in reasonable agreement with SPR (Supplementary figure S1) (Table 2).

**Table 2:**
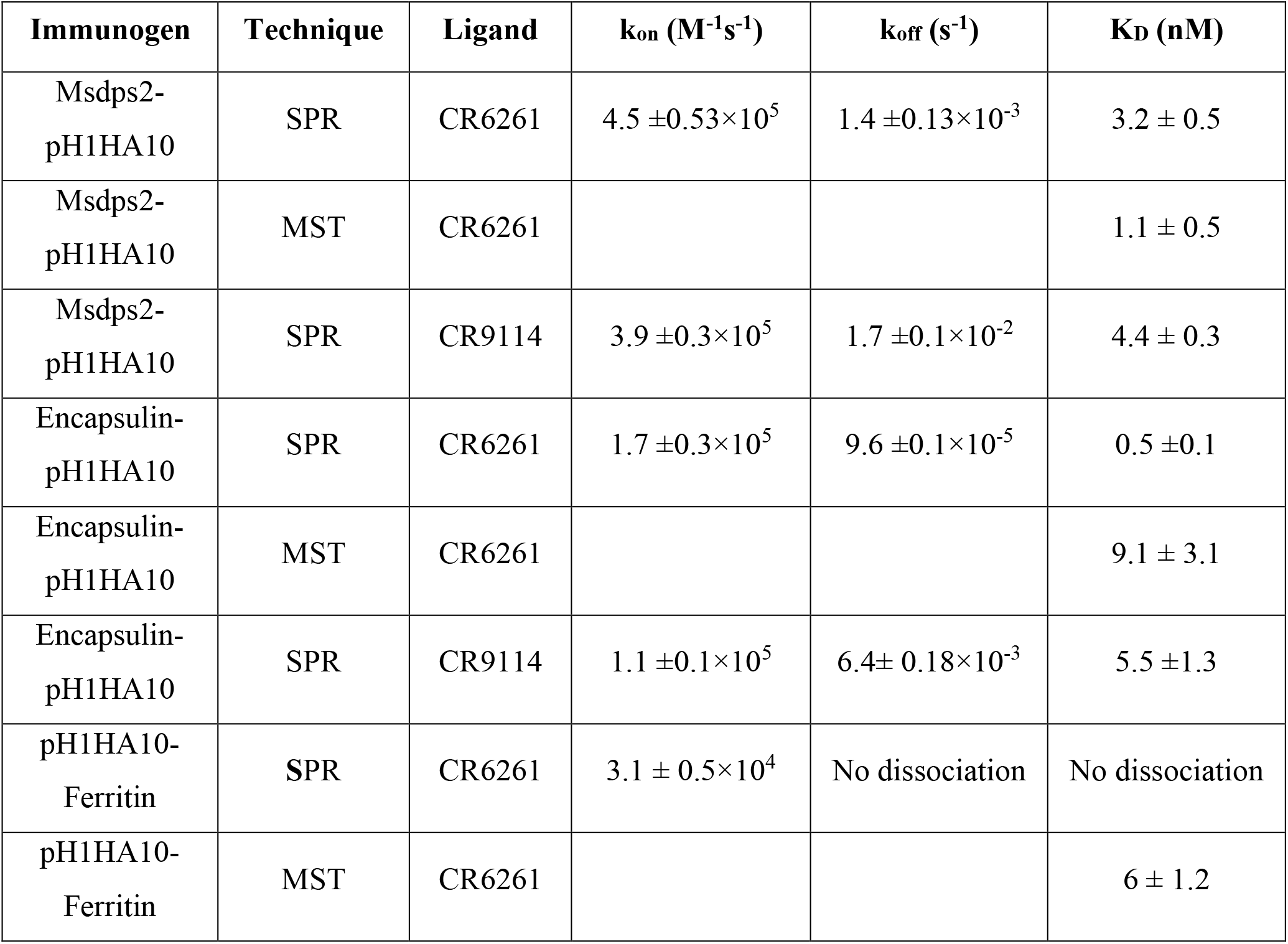
Binding parameters for interaction of nanoparticles with bNAb CR6261 measured by SPR and MST.

| **Immunogen** | **Technique** | **Ligand** | **$k_{on}$ ($M^{-1}s^{-1}$)** | **$k_{off}$ ($s^{-1}$)** | **$K_D$ (nM)** |
| --- | --- | --- | --- | --- | --- |
| Msdps2-pH1HA10 | SPR | CR6261 | $4.5 \pm 0.53 \times 10^5$ | $1.4 \pm 0.13 \times 10^{-3}$ | $3.2 \pm 0.5$ |
| Msdps2-pH1HA10 | MST | CR6261 | | | $1.1 \pm 0.5$ |
| Msdps2-pH1HA10 | SPR | CR9114 | $3.9 \pm 0.3 \times 10^5$ | $1.7 \pm 0.1 \times 10^{-2}$ | $4.4 \pm 0.3$ |
| Encapsulin-pH1HA10 | SPR | CR6261 | $1.7 \pm 0.3 \times 10^5$ | $9.6 \pm 0.1 \times 10^{-5}$ | $0.5 \pm 0.1$ |
| Encapsulin-pH1HA10 | MST | CR6261 | | | $9.1 \pm 3.1$ |
| Encapsulin-pH1HA10 | SPR | CR9114 | $1.1 \pm 0.1 \times 10^5$ | $6.4 \pm 0.18 \times 10^{-3}$ | $5.5 \pm 1.3$ |
| pH1HA10-Ferritin | SPR | CR6261 | $3.1 \pm 0.5 \times 10^4$ | No dissociation | No dissociation |
| pH1HA10-Ferritin | MST | CR6261 | | | $6 \pm 1.2$ |

Thermal tolerance is a desirable feature of vaccine candidates, especially in low resource settings. MsDps2-pH1HA10 was moderately stable (T_m_: 55 °C), comparable to the soluble trimeric stem immunogen (T_m_: 63.4 °C (Supplementary figure S2)) and retained functional epitopes after transient exposure to 40°C for 60 mins. Encapsulin-pH1HA10 did not show a clear transition in nanoDSF, hence we followed its thermal unfolding through CD. Encapsulin-pH1HA10 displayed higher stability (T_m_: 67°C) and remained functional even after being subjected to thermal stress with temperatures as high as 80 °C for 60 mins. pH1HA10-Ferritin was comparable in stability to MsDsp2-pH1HA10 (T_m_: 51.9 °C) (Figures 3F-I).

### Structural elucidation of designed MsDps2-pH1HA10 nanoparticles

MsDps2 is a member of the Ferritin superfamily that comprises of multi-subunit cage like proteins with a hollow interior and bears structural resemblance to ferritin (54). The above data demonstrate that MsDps2 nanoparticles can successfully act as a multivalent immunogen display platform, able to scaffold a trimeric pH1HA10 immunogen close to the three-fold axis symmetry on the MsDps2 core. The pH1HA10 immunogen displayed on the MsDps2 core was undetectable in the negative staining TEM images and class averages due to the flexible nature of linker. To detect the pH1HA10 immunogen on the MsDps2 nanoparticle core, we carried out cryo-EM. The micrographs and reference free 2D classifications indicate that the MsDps2-pH1HA10 particles are monodispersed and pH1HA10 immunogen was detectable in many 2D class averages (Supplementary figure 3A-B). These results motivated us to determine the cryo-EM structure of the MsDps2-pH1HA10 nanoparticle. Initially, 2-3 rounds of reference free 2D classification were performed on the cryo-EM dataset to remove disordered, beam damaged particles and ice contamination. A clean dataset was used to perform 3D classification of MsDps2-pH1HA10 using a 220 Å particle diameter to separate the various conformations of the pH1HA10 extension. A spherical ball-like initial model was used without any extension for 3D classification. Interestingly, all the classes in 3D classification show four large densities sticking out from a ball-like core particle. These four large densities appear at the same position as the three-fold axis symmetry of the core MsDps2 particle. Thus, these four large, elongated densities are pH1HA10 extensions, connected with the ball-like MsDps2 core structure (Supplementary figure 3C). This indicates that the majority of the MsDps2 core subunits are attached with pH1HA10. However, some classes (class 01, class 02, and class 03) show a weak connection between MsDps2 core and pH1HA10. These weak connections between core and pH1HA10 are likely due to the inherent flexibility of the linker connecting the MsDps2 core and pH1HA10.

Stable classes were selected for 3D reconstruction, and the 3D structure was calculated at a resolution of 8.5 Å. Overall structural features of the final 3D reconstruction are similar with stable 3D classes (class 04, class 05, class 06, class 07, and class 08), where four clear extensions at the three-fold axis symmetry of the MsDps2 core are visible (Figure 4A). Furthermore, the available crystal structure of MsDps2 core (PDB id-2Z90) and computationally generated pH1HA10 structure was manually fit into the cryo-EM density map, and the overall fitting of the atomic model strongly supports the moderate-resolution cryo-EM map. Further, the modelled dimensions of the pH1HA10 extension and available MsDps2 crystal structure sum up to the dimensions of the 3-D cryo-EM map of MsDps2-pH1HA10 thus verifying the design (Figure 4B-F). Due to the internal flexibility of the flexible linker between pH1HA10 and MsDps2, we are unable to resolve a near-atomic resolution structure of the MsDps2-pH1HA10 nanoparticles. In summary, the Mspdp2-pH1HA10 nanoparticle with four stable extensions is characterized at 8.5 Å showing 4 trimeric stems displayed on the MsDps2 core.

**Figure 4:**
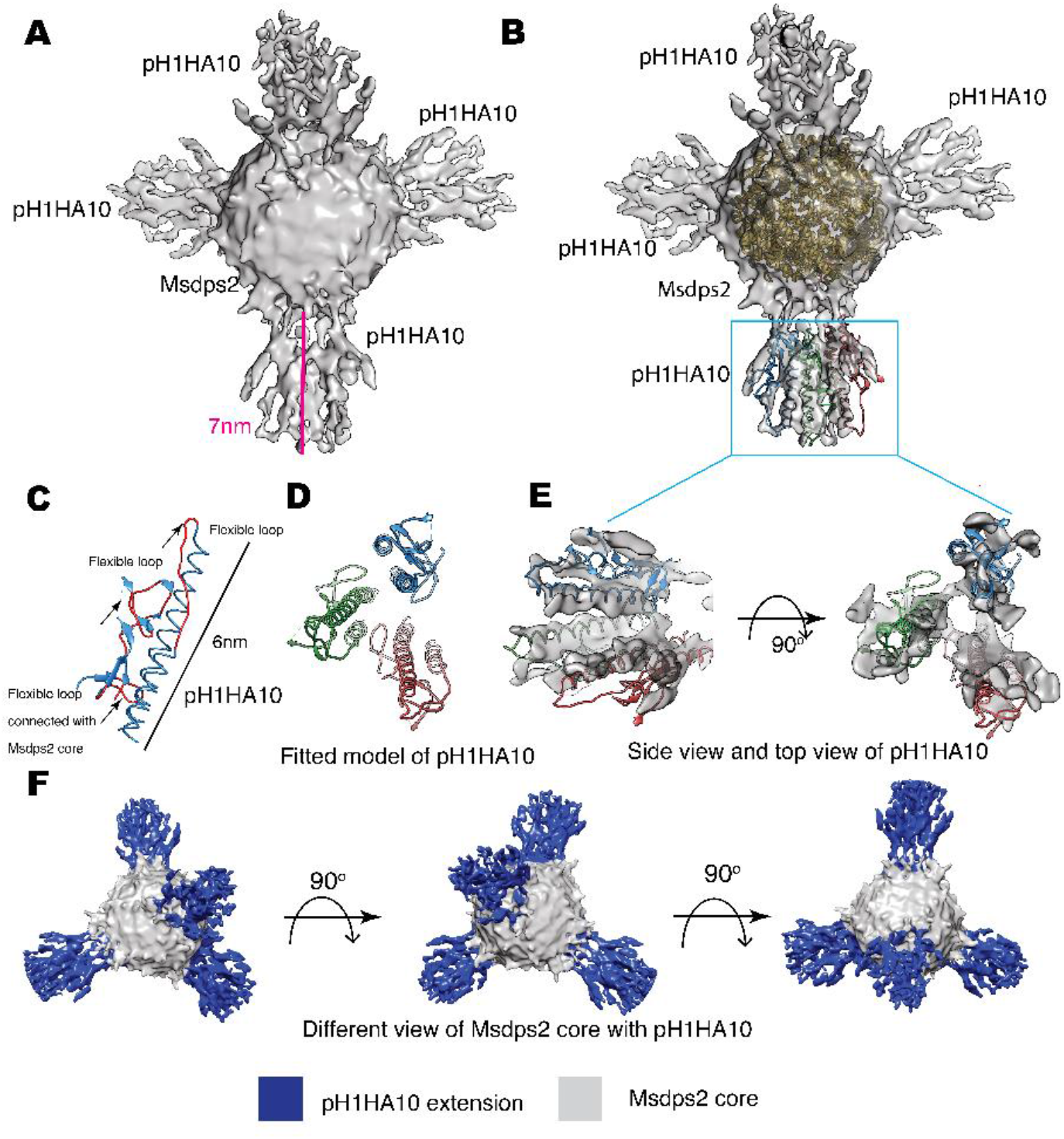
Cryo-EM structure of MsDps2-pH1HA10: **(A)** Solid representation of cryo-EM reconstruction of MsDps2-pH1HA10 shows four pH1HA10 extensions from MsDps2 core. Length of the pH1HA10 with linker is highlighted with a pink scale bar and length is 7 nm. **(B)** MsDps2 crystal structure (2Z90) is docked into the transparent view of the core region of MsDps2-pH1HA10. The pH1HA10 trimeric structure is fitted into the extended region of the cryo-EM map. **(C)** Modelled structure of pH1HA10 protomer. Flexible loops are shown in red. Dimension of pH1HA10 model structure is calculated to be 6 nm. **(D)** Top view of modelled structure of pH1HA10 trimer. **(E)** Side and top view of the transparent cryo-EM map of pH1HA10 trimeric extension. The protomers in pH1HA10 trimers are shown in blue, red and green respectively. **(F)** Different views of refined 3-D cryo-EM map of MsDps2-pH1HA10. pH1HA10 extension is shown in blue and MsDps2 core is shown in grey.

### Mice immunization and challenge studies

We examined the immunogenicity of SWE adjuvanted stem-nanoparticles in BALB/c mice. Animals were immunized intramuscularly at days 0 and 21 with 20 μg doses of the adjuvanted antigen. H1HA10-Foldon was immunized as a trimeric control. All proteins were adjuvanted with the MF59 equivalent oil in water emulsion, SWE (55). We employed this modality because intramuscular immunization is widely used in mass vaccinations, MF59 has an extensive safety record in humans and SWE is publicly available in GMP grade. One group of mice was immunized with PBS and acted as an unvaccinated control. Stem binding titers were evaluated at two time points, 2 weeks post prime (Wk2) and 2 weeks post boost (Wk5) through ELISA, for binding antibodies against the displayed trimeric stem, pH1HA10-Iz as the antigen. Iz is a synthetic trimerization domain (29). Use of this trimeric stem fusion makes it possible to accurately measure stem specific titers independent of the presence of foldon specific antibodies. Surprisingly, the nanoparticle elicited stem binding titers were not significantly higher than those of the trimer at either time point. This did not translate into a reduced protective efficacy, as complete protection was observed in case of nanoparticles, except for pH1HA10-Ferritin. MsDps2-pH1HA10 and pH1HA10-Ferritin elicited 2-fold and 14-fold lower stem binding titers, respectively, compared to the soluble trimer, pH1HA10-Foldon (MsDps2-pH1HA10 GMT: 14703, pH1HA10-Ferritin GMT: 2111, pH1HA10-Foldon GMT: 29407). Encapsulin-pH1HA10, which displayed 180 copies of the monomeric pH1HA10 elicited 4-fold lower stem binding antibodies (Encapsulin-pH1HA10 GMT: 7352) (Figure 5A). Post boost, binding titers against the ectodomain were also probed. For this, foldon was removed from the ectodomain-foldon protein through protease cleavage so as to eliminate foldon specific titers. In general, ectodomain binding titers were 2.3 to 5.3-fold lower compared to the pH1HA10-Iz titers. Encapsulin-pH1HA10 elicited the highest ectodomain binding titers among all the immunogens tested, whereas pH1HA10-Ferritin elicited the lowest binding titers (MsDps2-pH1HA10 GMT: 6400, Encapsulin-pH1HA10 GMT: 7352, pH1HA10-Ferritin GMT: 400, pH1HA10-Foldon GMT: 3676) (Figure 5B). Nanoparticle displayed antigens can also elicit antibodies against the scaffold platform. We observed scaffold titers as follows-MsDps2 GMT: 22286, Ferritin: 1393 and Encapsulin GMT: 16890 (Figure 5C). Microneutralization titers elicited by the nanoparticle immunogens correlate with the stem and ectodomain binding titers and approximately parallel the observed protection (Bel09 virus-MsDps2-pH1HA10 GMT: 65, Encapsulin-pH1HA10 GMT: 43, pH1HA10-Ferritin GMT: 20, pH1HA10-Foldon GMT: 49; PR8 virus-MsDps2-pH1HA10 GMT: 99, Encapsulin-pH1HA10 GMT: 70, pH1HA10-Ferritin GMT: 22, pH1HA10-Foldon GMT: 86) (Figure 5D-E). We also assessed the sera’s ability to compete with stem directed, broadly neutralizing mAB, CR6261. All the nanoparticle displayed immunogens competed with CR6261 with modest titers (MsDps2-pH1HA10 IC_50_: 511, Encapsulin-pH1HA10 IC_50_: 1746, pH1HA10-Ferritin IC_50_: 240, pH1HA10-Foldon IC_50_: 1290) (Figure 5F).

**Figure 5:**
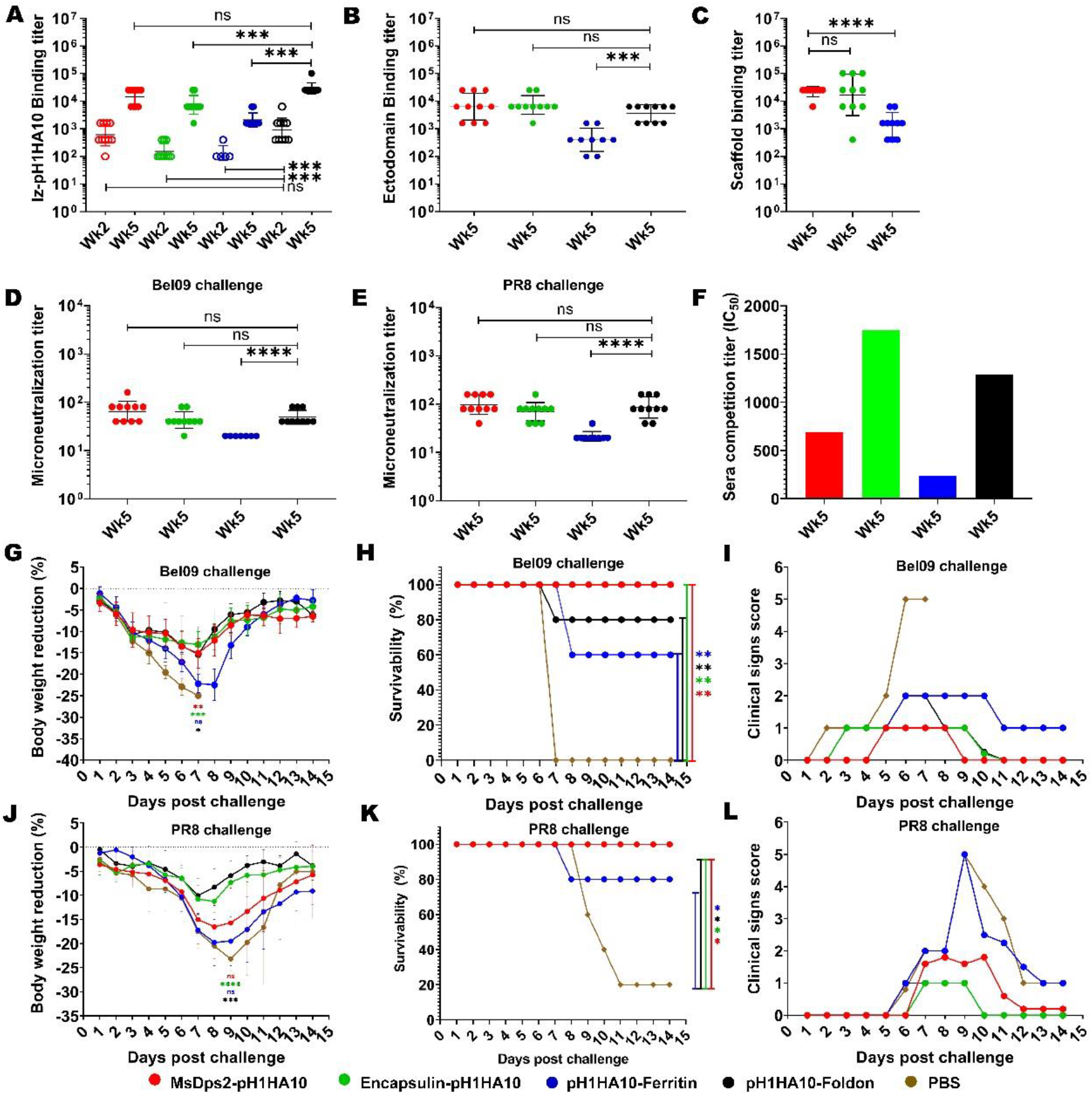
Mice immunization and challenge studies. Mice (n=5) were intramuscularly immunized with stem-nanoparticle immunogens-MsDps2-pH1HA10, Encapsulin-pH1HA10 and pH1HA10-Ferritin on days 0 and 21 with 20μg of SWE adjuvanted antigen. SWE adjuvanted pH1HA10-Foldon and PBS immunized groups acted as trimeric antigen vaccinated and unvaccinated controls, respectively. Sera was collected at days 14 (week 2) and 35 (week 5) and examined for binding titers against **(A)** pH1HA10-Iz and **(B)** Ectodomain (Ectodomain binding titers at week 5 are shown.). Scaffold titers at week 5 are shown in **(C)**. Comparison of ELISA binding titer between the pH1HA10-Foldon immunized and nanoparticle displayed pH1HA10 immunized groups was done using two-tailed Mann-Whitney t test. (ns indicates not significant, * indicates p < 0.05, ** indicates p < 0.01, and **** indicates p < 0.0001). Microneutralization titers were evaluated at week 5 to assess neutralizing antibodies against **(D)** Bel09 and **(E)** PR8. Groups were compared using two-tailed Mann Whitney t test (ns indicates not significant, * indicates p < 0.05, ** indicates p < 0.01, and **** indicates p < 0.0001). **(F)** Competition assay of pooled sera of each antigen with CR6261 for binding to immobilized pH1HA10-Foldon through SPR. Post immunization, mice were challenged with either 10423 pfu (10MLD_50_) of vaccine matched, H1N1 A/Belgium/145-MA/2009 (Bel09) virus **(G-H)**, or 270 pfu (10MLD_50_) of vaccine mismatched, H1N1 A/Puerto Rico/8/1934 (PR8) strain **(J-K)**. Mice morbidity **(G, J)** and mortality **(H, K)** health and weight change was monitored up to 2 weeks post challenge. Survivability between vaccinated and unvaccinated groups was compared using a two-way ANOVA test with Bonferroni correction (p < 0.01 (*), p < 0.001 (**), p < 0.0001 (****)). Multiple T test with Bonferroni Dunn correction was used to compare body weight reduction between groups (p < 0.01 (*), p < 0.001 (**), p < 0.0001 (****)). **(F, I)** Individual group averaged clinical scores are shown after **(I)** Bel09 or **(L)** PR8 challenge.

Following immunization, mice (n=5) were intranasally challenged with a lethal dose (10MLD_50_) of vaccine matched H1N1 A/Belgium/145-MA/2009 replicative virus. pH1HA10-Foldon immunized, and PBS immunized animal groups acted as trimeric vaccinated and unvaccinated controls, respectively. Post infection, vaccinated groups showed some weight loss, but regained body weight 8 days after challenge, in contrast to the unvaccinated group (Figure 5G). MsDps2-pH1HA10 and Encapsulin-pH1HA10 protected completely against a lethal homologous challenge, whereas three animals in the pH1HA10-Ferritin vaccinated group, one animal in the trimeric immunogen vaccinated group and all the animals in the unvaccinated group succumbed to the infection, respectively (Figure 5H). In general, MsDps2-pH1HA10 and Encapsulin-pH1HA10 immunized mice showed lower clinical signs compared to pH1HA10-Ferritin which disappeared on day 9 and day 10, respectively. pH1HA10-Ferritin showed moderate clinical symptoms throughout the observation period. pH1HA10-Foldon immunized group showed mild clinical signs which disappeared on day 11. In contrast, unvaccinated group showed high clinical signs until day 7 after which all the animals in the group succumbed to the infection (Figure 5I). We, next, examined the protective efficacy of the stem immunogens against a highly divergent H1N1 A/Puerto Rico/8/1934 replicative virus. With the exception of pH1HA10-Foldon and Encapsulin-pH1HA10 immunized animals, greater body weight reduction was observed compared to the homologous challenge, however, all surviving animals regained weight post clearance of infection (Figure 5J). MsDps2-pH1HA10, Encapsulin-pH1HA10 and pH1HA10-Foldon vaccinated groups were completely protected against the heterologous challenge, whereas pH1HA10-Ferritin afforded partial protection. In contrast, only one animal survived in the unvaccinated control group (Figure 5K). Encapsulin-pH1HA10 and pH1HA10-Foldon showed the lowest clinical signs which disappeared on day 10, whereas MsDps2-pH1HA10 showed some symptoms until day 12. In contrast, pH1HA10-Ferritin and PBS immunized groups showed very high clinical scores which persisted until day 13 (Figure 5I). In summary, all the tested antigens elicited moderate titers of binding antibodies and protected completely or partially against lethal homologous and heterologous challenge. Specifically, a small but statistically significant enhancement in protection was observed in case of the nanoparticle displayed stem immunogens over the soluble trimer in the homologous challenge, whereas both nanoparticle displayed stem immunogens and trimeric stem immunogen protected mice against a heterologous challenge with similar efficacy. Encapsulin-pH1HA10 showed the lowest body weight reduction among all the tested immunogens in the homologous challenge, whereas pH1HA10-Foldon showed the lowest body weight reduction in the heterologous challenge.

## Discussion

Focussing the immune response to conserved epitopes on HA is a promising strategy towards the development of ‘universal vaccine candidates (21,56,57). This has resulted in the development of various subunit vaccine designs eliciting responses to HA head or HA stem (24,29,58–61).

Several recent studies have described self-assembling protein nanoparticles that induce broadly neutralizing antibody responses against influenza virus in mice and ferrets (23,24). Using similar approaches to ours, immunogens derived from the surface proteins of HIV-1, and RSV have been displayed on self-assembling, naturally occurring nanoparticles, as well as computationally designed nanoparticle scaffolds (62,63). Recently, there was a report of a double-layered nanoparticle that induced broad protection against divergent influenza strains (64). In recent Covid-19 pandemic scenario, several groups have successfully utilized protein nanoparticle platforms to display SARS-COV2 antigens as measure to combat the deadly virus (65–69).

Owing to the flexible nature of the displayed immunogens, it has not been possible to obtain a high resolution cryo-EM map for any nanoparticle immunogen fusion to date. However, in all the above cases, one or more components were expressed in mammalian cell culture. In contrast, in the present case, the immunogen is bacterially expressed; which has several advantages in terms of cost and scalability.

In this work, we have successfully shown the design, biophysical characterization, and immunogenicity of bacterially expressed, different sized nanoparticle fusion constructs where the stem domain subunit immunogen pH1HA10 was displayed on 12-mer MsDps2, 24-mer Ferritin and 180-mer Encapsulin nanoparticles. The nanoparticle fusion proteins were purified in moderate to good yield and were properly folded as assessed by their ability to bind stem specific conformational antibodies. Size exclusion analysis and negative stain EM data clearly indicate proper homogenous particle formation for both Msdps2 and Ferritin fused immunogens. On the other hand, the Encapsulin fusion construct exhibited a heterogenous oligomeric particle distribution.

Since the immunogen extensions were not visible in negative stain images, we employed Cryo-EM to detect the antigen displayed on the nanoparticle platform. Several labs have previously carried out cryo-EM analysis of nanoparticle immunogen display platforms since a high resolution cryo-EM map provides structural information on the displayed immunogens. The Cryo-EM map of the HA stem displayed on Ferritin nanoparticles was previously obtained at a low resolution of 16 Å (24). In 2019, the Cryo-EM structure of a computationally designed nanoparticle I53 displaying the Respiratory Syncytial Virus (RSV) virus was solved at a resolution of 6.3 Å (63). Recently, there are reports of cryo-EM maps of several computationally designed two-component protein nanoparticle immunogen display platforms at a resolution of 3.9-6.67 Å (70,71). In most of these cryo-EM studies, the displayed immunogens were more poorly resolved than the corresponding nanoparticle core. Here, we obtained a moderate resolution cryo-EM map of a naturally occurring protein nanoparticle immunogen display platform (MsDps2 displaying influenza stem immunogen). The immunogens (pH1HA10) in the Cryo-EM maps were displayed on the three-fold symmetry axes on the nanoparticle MsDps2. The immunogens are attached to the MsDps2 nanoparticle core by 9 amino acid linkers (GSAGSAGAS). GS (Gly Ser) linkers are one of the most widely used flexible linkers in fusion protein designs owing to their small size and hydrophobic character. Further, these linkers can form hydrogen bonds in an aqueous solution thus avoiding undesirable interactions with the displayed protein (72,73). A plausible reason for the moderate cryo-EM resolution obtained in the study is this flexible nature of the displayed immunogen which may lead to distortion of global alignment thus hampering the overall resolution of the MsDps2-pH1HA10 fusion protein. Alternative linker designs to decrease the flexibility of the displayed antigen might result in a higher resolution structure, though whether this will enhance immunogenicity in unclear. Several previous studies have indicated that flexibility of the particle does not depend on the beam-induced motion or mechanical drift. An increase in flexibility leads to relative movement of different regions of the molecule which reduces the quality of the final data with a single prolonged exposure (74). Our result is in accord with the previous cryo-EM studies of nanoparticle immunogen fusion by other labs where the 3-D reconstruction of complete nanoparticles gave rise to highly diffused and poorly resolved displayed immunogen as compared to the nanoparticle core (63,71). Focused refinement of the MsDps2 core gave rise to a high resolution cryo-EM map of the core with a resolution of 3.45 Å. This enhancement of resolution upon focused refinement confirms our hypothesis that linker flexibility is responsible for the loss of resolution in the overall cryo-EM map. The available crystal structure of MsDps2 (PDB) was found to fit very well to the Cryo-EM map of the MsDps2 core. The whole Cryo-EM map generation was *de-novo* without considering any available crystal structures as a template.

Intramuscular immunizations of mice with the SWE adjuvanted nanoparticle fusion constructs indicated that the nanoparticle displayed antigens were immunogenic, though surprisingly there was no substantial enhancement over described parent trimeric stem immunogen (29). Increase in the multimeric display valency from 12-mer MsDps2 to 180-mer Encapsulin did not lead to any increase in the antibody titers likely due to steric repulsion between adjacent displayed immunogens in Encapsulin-pH1HA10 leading to hindered BCR interaction with the available epitopes. However, it did show qualitatively superior performance in terms of lower morbidity, relative to the other nanoparticles tested. When challenged with a lethal dose of homologous influenza strain, (A/Belgium/145-MA/2009), mice immunized with the nanoparticle fusion constructs were completely protected, except pH1HA10-Ferritin which showed partial protection. Although the vaccinated group showed substantial weight loss initially, all the mice regained their body weight shortly. In a heterologous challenge study with a lethal dose of a divergent PR8 strain, Msdps2-pH1HA10 and Encapsulin-pH1HA10 immunized mice were fully protected while pH1HA10-Ferritin immunized mice were partially protected. Soluble trimer showed partial protection in the homologous challenge, whereas complete protection in the heterologous challenge. In the PBS control group, even though a 10MLD_50_ dose was used, 1 of 5 mice survived the PR8 challenge while all the mice in the Bel09 challenged group died. The MLD_50_ determined by us for PR8 is 27 pfu which is in line with published values of 25 pfu (75) and showed 100% mortality during MLD_50_ estimation. It is possible that the dose administered to the PR8 group in the present study was less than 10MLD_50_ because of loss of infectivity upon −80°C storage. Multivalent display facilitated by nanoparticles effectively increases the number of antigens presented to the host immune system, while at the same time ensuing an avidity response sparked by multivalent engagement of B-cells. This could allow for dose sparing strategies, especially beneficial in low resource settings. We will examine relative immunogenicity and protection at lower doses in a future study. In summary, the study discusses the design of various sized, bacterially expressed, nanoparticle display platforms which were properly folded and showed high affinity binding to conformation specific stem based bnAbs. Using intramuscular immunization with SWE adjuvanted formulations, the nanoparticle displayed antigens elicited moderate titers of binding antibodies and protected completely or partially against lethal homologous and heterologous challenge. This route of administration is suitable for mass immunizations, and the adjuvant is publicly available in GMP grade. Overall, Encapsulin-pH1HA10 showed the best protective immune response in terms of mice mortality and morbidity after homologous and heterologous challenge. In conclusion, all the vaccinated groups conferred protection compared to the unvaccinated group. A small but statistically significant improvement was observed in the protective efficacy of the nanoparticle displayed immunogens compared to the parent, soluble trimer in the homologous challenge. However, there was no such significant enhancement relative to the parent, soluble trimer in the heterologous challenge. This is agreement to an earlier study (69) where the nanoparticle displayed derivatives of the receptor binding domain of SARS-CoV-2 did not show any enhancement in immunogenicity relative to soluble trimer. Given the higher yields, ease of characterization and better manufacturability in the case of the present HA stem trimer, the soluble trimer is a preferred candidate immunogen relative to the nanoparticle displayed derivatives. Unlike most other designs where one or more components are expressed in mammalian cells, the present immunogens can be rapidly and cheaply produced in a pandemic, including in low-resource settings.

## Author Contributions

RV, UK, and SK designed the study. RV, UK, and SK analysed and interpreted most of the data. UK designed the constructs. UK and SK purified and characterized all nanoparticle fusion constructs. SK carried out the thermal stability, competition assays and statistical analyses. SD designed the cryo-EM data collection and image processing. PG and SD carried out all the negative stain EM and cryo-EM data processing, data analysis and data interpretation. AU did some of the protein purifications. ELISA were done by MD, PR and SK. Challenge studies were carried out by MB. Microneutralization assay was carried out by PR. SP formulated the immunogens and along with GC coordinated the animal immunizations. UK, SK, and RV wrote the manuscript with critical input from all authors. All authors contributed to the article and approved the submitted version.

## Acknowledgements

We thank Prof Dipankar Chatterji (Molecular Biophysics Unit, IISc) for providing us with the MsDps2 plasmid. We acknowledge Ms. Nidhi Mittal for providing the H1 Ectodomain protein for ELISA studies and Ms. Nisha Bhandari for providing the pH1AH10-Foldon protein for SPR and nanoDSF studies. We thank Mr. Gopinath Chattopadhyay for his help in carrying out competition assay through SPR. We acknowledge technical support from Ms. Nonavinakere Seetharam Srilatha for assistance during SPR experiments. We are thankful to Dr. Xavier Saelens, VIB-UGent Center for Medical Biology, Belgium, for kindly providing the H1N1 A/Belgium/145-MA/2009 viral strain and to Dr. Shashank Tripathi, Center for Infectious Disease Research, IISc, India, for providing the H1N1 A/Puerto Rico/8/1934 viral strain. We sincerely acknowledge funding from Uchchatar Avishkar Yojana (UAY), Ministry of Human Resource Development (IISc_005) grant to RV, DBT-ENDFLU (BT/IN/EU-INF/15/RV/19-20) funded by European Union’s Horizon 2020 research and innovation programme under grant agreement No. 874650 and the Department of Biotechnology, Ministry of Science and Technology, Government of India (BT/IN/EU-INF/15/RV/19-20), and a grant from the Bill and Melinda Gates Foundation (INV-005948). We acknowledge funding for infrastructural support from DST-FIST, UGC Centre of Advanced Study, MHRD-FAST, the DBT-IISc Partnership Program, and of a JC Bose Fellowship from DST to R.V. S.D. acknowledges the support of the DBT Cryo-EM facility (BT/INF/22/SP22844/2017SM). SK acknowledges fellowship from DBT-JRF (DBT/JRF/BET-18/I/2018/AL/168). UK acknowledges fellowship from UGC-CSIR.

## References

1. Reed C, Angulo FJ, Swerdlow DL, Lipsitch M, Meltzer MI, Jernigan D, et al. Estimates of the prevalence of pandemic (H1N1) 2009, United States, April-July 2009. Emerg Infect Dis [Internet]. 2009 Dec;15(12):2004–7. Available from: http://www.ncbi.nlm.nih.gov/pubmed/19961687

2. Tong S, Zhu X, Li Y, Shi M, Zhang J, Bourgeois M, et al. New World Bats Harbor Diverse Influenza A Viruses. PLoS Pathog. 2013;9(10).

3. Shi Y, Wu Y, Zhang W, Qi J, Gao GF. Enabling the “host jump”: structural determinants of receptor-binding specificity in influenza A viruses. Nat Rev Microbiol [Internet]. 2014 Dec;12(12):822–31. Available from: http://www.ncbi.nlm.nih.gov/pubmed/25383601

4. Girard MP, Cherian T, Pervikov Y, Kieny MP. A review of vaccine research and development: human acute respiratory infections. Vaccine [Internet]. 2005 Dec 30;23(50):5708–24. Available from: http://www.ncbi.nlm.nih.gov/pubmed/16154667

5. Belser JA, Lu X, Maines TR, Smith C, Li Y, Donis RO, et al. Pathogenesis of avian influenza (H7) virus infection in mice and ferrets: enhanced virulence of Eurasian H7N7 viruses isolated from humans. J Virol [Internet]. 2007 Oct;81(20):11139–47. Available from: http://www.ncbi.nlm.nih.gov/pubmed/17686867

6. Pica N, Palese P. Toward a universal influenza virus vaccine: prospects and challenges. Annu Rev Med [Internet]. 2013;64:189–202. Available from: http://www.ncbi.nlm.nih.gov/pubmed/23327522

7. Berlanda Scorza F, Tsvetnitsky V, Donnelly JJ. Universal influenza vaccines: Shifting to better vaccines. Vaccine [Internet]. 2016;34(26):2926–33. Available from: http://www.ncbi.nlm.nih.gov/pubmed/27038130

8. Paules CI, Marston HD, Eisinger RW, Baltimore D, Fauci AS. The Pathway to a Universal Influenza Vaccine. Immunity [Internet]. 2017;47(4):599–603. Available from: http://www.ncbi.nlm.nih.gov/pubmed/29045889

9. Safdar A, Cox MMJ. Baculovirus-expressed influenza vaccine. A novel technology for safe and expeditious vaccine production for human use. Expert Opin Investig Drugs [Internet]. 2007 Jul;16(7):927–34. Available from: http://www.ncbi.nlm.nih.gov/pubmed/17594180

10. Skehel JJ, Wiley DC. Receptor binding and membrane fusion in virus entry: the influenza hemagglutinin. Annu Rev Biochem [Internet]. 2000;69:531–69. Available from: http://www.ncbi.nlm.nih.gov/pubmed/10966468

11. Wiley DC, Wilson IA, Skehel JJ. Structural identification of the antibody-binding sites of Hong Kong influenza haemagglutinin and their involvement in antigenic variation. Nature [Internet]. 1981 Jan 29;289(5796):373–8. Available from: http://www.ncbi.nlm.nih.gov/pubmed/6162101

12. Both GW, Sleigh MJ, Cox NJ, Kendal AP. Antigenic drift in influenza virus H3 hemagglutinin from 1968 to 1980: multiple evolutionary pathways and sequential amino acid changes at key antigenic sites. J Virol [Internet]. 1983 Oct;48(1):52–60. Available from: http://www.ncbi.nlm.nih.gov/pubmed/6193288

13. Nobusawa E, Aoyama T, Kato H, Suzuki Y, Tateno Y, Nakajima K. Comparison of complete amino acid sequences and receptor-binding properties among 13 serotypes of hemagglutinins of influenza A viruses. Virology [Internet]. 1991 Jun;182(2):475–85. Available from: http://www.ncbi.nlm.nih.gov/pubmed/2024485

14. Corti D, Voss J, Gamblin SJ, Codoni G, Macagno A, Jarrossay D, et al. A neutralizing antibody selected from plasma cells that binds to group 1 and group 2 influenza A hemagglutinins. Science (80-). 2011;333(6044):850–6.

15. Ekiert DC, Bhabha G, Elsliger M-A, Friesen RHE, Jongeneelen M, Throsby M, et al. Antibody recognition of a highly conserved influenza virus epitope. Science [Internet]. 2009 Apr 10;324(5924):246–51. Available from: http://www.ncbi.nlm.nih.gov/pubmed/19251591

16. Krammer F, Palese P. Influenza virus hemagglutinin stalk-based antibodies and vaccines. Curr Opin Virol [Internet]. 2013 Oct;3(5):521–30. Available from: http://www.ncbi.nlm.nih.gov/pubmed/23978327

17. Ekiert DC, Friesen RHE, Bhabha G, Kwaks T, Jongeneelen M, Yu W, et al. A highly conserved neutralizing epitope on group 2 influenza A viruses. Science (80-). 2011;333(6044):843–50.

18. Okuno Y, Isegawa Y, Sasao F, Ueda S. A common neutralizing epitope conserved between the hemagglutinins of influenza A virus H1 and H2 strains. J Virol [Internet]. 1993 May;67(5):2552–8. Available from: http://www.ncbi.nlm.nih.gov/pubmed/7682624

19. Sanchez-Fauquier A, Guillen M, Martin J, Kendal AP, Melero JA. Conservation of epitopes recognized by monoclonal antibodies against the separated subunits of influenza hemagglutinin among type A viruses of the same and different subtypes. Arch Virol [Internet]. 1991;116(1–4):285–92. Available from: http://www.ncbi.nlm.nih.gov/pubmed/1705790

20. Throsby M, van den Brink E, Jongeneelen M, Poon LLM, Alard P, Cornelissen L, et al. Heterosubtypic neutralizing monoclonal antibodies cross-protective against H5N1 and H1N1 recovered from human IgM+ memory B cells. PLoS One [Internet]. 2008;3(12):e3942. Available from: http://www.ncbi.nlm.nih.gov/pubmed/19079604

21. Sui J, Hwang WC, Perez S, Wei G, Aird D, Chen L, et al. Structural and functional bases for broad-spectrum neutralization of avian and human influenza A viruses. Nat Struct Mol Biol [Internet]. 2009 Mar;16(3):265–73. Available from: http://www.ncbi.nlm.nih.gov/pubmed/19234466

22. Lee LA, Wang Q. Adaptations of nanoscale viruses and other protein cages for medical applications. Vol. 2, Nanomedicine: Nanotechnology, Biology, and Medicine. 2006. p. 137–49.

23. Kanekiyo M, Wei CJ, Yassine HM, McTamney PM, Boyington JC, Whittle JRR, et al. Self-assembling influenza nanoparticle vaccines elicit broadly neutralizing H1N1 antibodies. Nature. 2013;499(7456):102–6.

24. Yassine HM, Boyington JC, McTamney PM, Wei CJ, Kanekiyo M, Kong WP, et al. Hemagglutinin-stem nanoparticles generate heterosubtypic influenza protection. Nat Med. 2015;21(9):1065–70.

25. Georgiev IS, Joyce MG, Chen RE, Leung K, McKee K, Druz A, et al. Two-Component Ferritin Nanoparticles for Multimerization of Diverse Trimeric Antigens. ACS Infect Dis [Internet]. 2018;4(5):788–96. Available from: http://www.ncbi.nlm.nih.gov/pubmed/29451984

26. Corbett KS, Moin SM, Yassine HM, Cagigi A, Kanekiyo M, Boyoglu-Barnum S, et al. Design of Nanoparticulate Group 2 Influenza Virus Hemagglutinin Stem Antigens That Activate Unmutated Ancestor B Cell Receptors of Broadly Neutralizing Antibody Lineages. MBio [Internet]. 2019;10(1). Available from: http://www.ncbi.nlm.nih.gov/pubmed/30808695

27. Roy S, Saraswathi R, Chatterji D, Vijayan M. Structural Studies on the Second Mycobacterium smegmatis Dps: Invariant and Variable Features of Structure, Assembly and Function. J Mol Biol. 2008;375(4):948–59.

28. McHugh CA, Fontana J, Nemecek D, Cheng N, Aksyuk AA, Heymann JB, et al. A virus capsid-like nanocompartment that stores iron and protects bacteria from oxidative stress. EMBO J [Internet]. 2014 Sep 1;33(17):1896–911. Available from: http://www.ncbi.nlm.nih.gov/pubmed/25024436

29. Mallajosyula VVA, Citron M, Ferrara F, Lu X, Callahan C, Heidecker GJ, et al. Influenza hemagglutinin stem-fragment immunogen elicits broadly neutralizing antibodies and confers heterologous protection. Proc Natl Acad Sci U S A. 2014;111(25).

30. Bommakanti G, Lu X, Citron MP, Najar TA, Heidecker GJ, ter Meulen J, et al. Design of Escherichia coli-Expressed Stalk Domain Immunogens of H1N1 Hemagglutinin That Protect Mice from Lethal Challenge. J Virol. 2012;86(24):13434–44.

31. Sharma A, Chattopadhyay G, Chopra P, Bhasin M, Thakur C, Agarwal S, et al. VapC21 Toxin Contributes to Drug-Tolerance and Interacts With Non-cognate VapB32 Antitoxin in Mycobacterium tuberculosis. Front Microbiol [Internet]. 2020;11:2037. Available from: http://www.ncbi.nlm.nih.gov/pubmed/33042034

32. Jerabek-Willemsen M, Wienken CJ, Braun D, Baaske P, Duhr S. Molecular interaction studies using microscale thermophoresis. Assay Drug Dev Technol [Internet]. 2011 Aug;9(4):342–53. Available from: http://www.ncbi.nlm.nih.gov/pubmed/21812660

33. Jerabek-Willemsen M, André T, Wanner R, Roth HM, Duhr S, Baaske P, et al. MicroScale Thermophoresis: Interaction analysis and beyond. J Mol Struct [Internet]. 2014 Dec;1077:101–13. Available from: https://linkinghub.elsevier.com/retrieve/pii/S0022286014002750

34. Wienken CJ, Baaske P, Rothbauer U, Braun D, Duhr S. Protein-binding assays in biological liquids using microscale thermophoresis. Nat Commun [Internet]. 2010 Oct 19;1:100. Available from: http://www.ncbi.nlm.nih.gov/pubmed/20981028

35. Chattopadhyay G, Varadarajan R. Facile measurement of protein stability and folding kinetics using a nano differential scanning fluorimeter. Protein Sci [Internet]. 2019;28(6):1127–34. Available from: http://www.ncbi.nlm.nih.gov/pubmed/30993730

36. Kumar S, Panda H, Makhdoomi MA, Mishra N, Safdari HA, Chawla H, et al. An HIV-1 Broadly Neutralizing Antibody from a Clade C-Infected Pediatric Elite Neutralizer Potently Neutralizes the Contemporaneous and Autologous Evolving Viruses. J Virol [Internet]. 2019;93(4). Available from: http://www.ncbi.nlm.nih.gov/pubmed/30429339

37. Ghosh E, Dwivedi H, Baidya M, Srivastava A, Kumari P, Stepniewski T, et al. Conformational Sensors and Domain Swapping Reveal Structural and Functional Differences between β-Arrestin Isoforms. Cell Rep [Internet]. 2019;28(13):3287–3299.e6. Available from: http://www.ncbi.nlm.nih.gov/pubmed/31553900

38. Bell JM, Chen M, Baldwin PR, Ludtke SJ. High resolution single particle refinement in EMAN2.1. Methods [Internet]. 2016;100:25–34. Available from: http://www.ncbi.nlm.nih.gov/pubmed/26931650

39. Zheng SQ, Palovcak E, Armache J-P, Verba KA, Cheng Y, Agard DA. MotionCor2: anisotropic correction of beam-induced motion for improved cryo-electron microscopy. Nat Methods [Internet]. 2017;14(4):331–2. Available from: http://www.ncbi.nlm.nih.gov/pubmed/28250466

40. Scheres SHW. RELION: implementation of a Bayesian approach to cryo-EM structure determination. J Struct Biol [Internet]. 2012 Dec;180(3):519–30. Available from: http://www.ncbi.nlm.nih.gov/pubmed/23000701

41. Kimanius D, Forsberg BO, Scheres SH, Lindahl E. Accelerated cryo-EM structure determination with parallelisation using GPUs in RELION-2. Elife [Internet]. 2016;5. Available from: http://www.ncbi.nlm.nih.gov/pubmed/27845625

42. Rohou A, Grigorieff N. CTFFIND4: Fast and accurate defocus estimation from electron micrographs. J Struct Biol [Internet]. 2015 Nov;192(2):216–21. Available from: http://www.ncbi.nlm.nih.gov/pubmed/26278980

43. van Heel M, Schatz M. Fourier shell correlation threshold criteria. J Struct Biol [Internet]. 2005 Sep;151(3):250–62. Available from: http://www.ncbi.nlm.nih.gov/pubmed/16125414

44. Pettersen EF, Goddard TD, Huang CC, Couch GS, Greenblatt DM, Meng EC, et al. UCSF Chimera--a visualization system for exploratory research and analysis. J Comput Chem [Internet]. 2004 Oct;25(13):1605–12. Available from: http://www.ncbi.nlm.nih.gov/pubmed/15264254

45. Xu R, Ekiert DC, Krause JC, Hai R, Crowe JE, Wilson IA. Structural basis of preexisting immunity to the 2009 H1N1 pandemic influenza virus. Science [Internet]. 2010 Apr 16;328(5976):357–60. Available from: http://www.ncbi.nlm.nih.gov/pubmed/20339031

46. Mittal N, Sengupta N, Malladi SK, Reddy P, Bhat M, Rajmani RS, et al. Protective Efficacy of Recombinant Influenza Hemagglutinin Ectodomain Fusions. Viruses [Internet]. 2021 Aug 27;13(9):1710. Available from: https://www.mdpi.com/1999-4915/13/9/1710

47. Nguyen B, Tolia NH. Protein-based antigen presentation platforms for nanoparticle vaccines. NPJ vaccines [Internet]. 2021 May 13;6(1):70. Available from: http://www.ncbi.nlm.nih.gov/pubmed/33986287

48. Pati R, Shevtsov M, Sonawane A. Nanoparticle Vaccines Against Infectious Diseases. Front Immunol [Internet]. 2018;9:2224. Available from: http://www.ncbi.nlm.nih.gov/pubmed/30337923

49. He L, de Val N, Morris CD, Vora N, Thinnes TC, Kong L, et al. Presenting native-like trimeric HIV-1 antigens with self-assembling nanoparticles. Nat Commun [Internet]. 2016;7:12041. Available from: http://www.ncbi.nlm.nih.gov/pubmed/27349934

50. Boyoglu-Barnum S, Ellis D, Gillespie RA, Hutchinson GB, Park Y-J, Moin SM, et al. Quadrivalent influenza nanoparticle vaccines induce broad protection. Nature [Internet]. 2021;592(7855):623–8. Available from: http://www.ncbi.nlm.nih.gov/pubmed/33762730

51. López-Sagaseta J, Malito E, Rappuoli R, Bottomley MJ. Self-assembling protein nanoparticles in the design of vaccines. Comput Struct Biotechnol J [Internet]. 2016;14:58–68. Available from: http://www.ncbi.nlm.nih.gov/pubmed/26862374

52. Bachmann MF, Rohrer UH, Kündig TM, Bürki K, Hengartner H, Zinkernagel RM. The influence of antigen organization on B cell responsiveness. Science [Internet]. 1993 Nov 26;262(5138):1448–51. Available from: http://www.ncbi.nlm.nih.gov/pubmed/8248784

53. Zhao L, Seth A, Wibowo N, Zhao C-X, Mitter N, Yu C, et al. Nanoparticle vaccines. Vaccine [Internet]. 2014 Jan 9;32(3):327–37. Available from: http://www.ncbi.nlm.nih.gov/pubmed/24295808

54. Zhang Y, Orner BP. Self-assembly in the ferritin nano-cage protein superfamily. Int J Mol Sci [Internet]. 2011;12(8):5406–21. Available from: http://www.ncbi.nlm.nih.gov/pubmed/21954367

55. O’Hagan DT, Ott GS, Nest G Van, Rappuoli R, Giudice G Del. The history of MF59(®) adjuvant: a phoenix that arose from the ashes. Expert Rev Vaccines [Internet]. 2013 Jan;12(1):13–30. Available from: http://www.ncbi.nlm.nih.gov/pubmed/23256736

56. Du L, Zhou Y, Jiang S. Research and development of universal influenza vaccines. Microbes Infect [Internet]. 2010 Apr;12(4):280–6. Available from: http://www.ncbi.nlm.nih.gov/pubmed/20079871

57. Steel J, Lowen AC, Wang TT, Yondola M, Gao Q, Haye K, et al. Influenza virus vaccine based on the conserved hemagglutinin stalk domain. MBio. 2010;1(1).

58. Impagliazzo A, Milder F, Kuipers H, Wagner M V., Zhu X, Hoffman RMB, et al. A stable trimeric influenza hemagglutinin stem as a broadly protective immunogen. Science (80-). 2015;349(6254):1301–6.

59. Sutton TC, Chakraborty S, Mallajosyula VVA, Lamirande EW, Ganti K, Bock KW, et al. Protective efficacy of influenza group 2 hemagglutinin stem-fragment immunogen vaccines. npj Vaccines. 2017;2(1).

60. Lu Y, Welsh JP, Swartz JR. Production and stabilization of the trimeric influenza hemagglutinin stem domain for potentially broadly protective influenza vaccines. Proc Natl Acad Sci U S A [Internet]. 2014 Jan 7;111(1):125–30. Available from: http://www.ncbi.nlm.nih.gov/pubmed/24344259

61. Xu R, Krause JC, McBride R, Paulson JC, Crowe JE, Wilson IA. A recurring motif for antibody recognition of the receptor-binding site of influenza hemagglutinin. Nat Struct Mol Biol [Internet]. 2013 Mar;20(3):363–70. Available from: http://www.ncbi.nlm.nih.gov/pubmed/23396351

62. Jardine J, Julien J-P, Menis S, Ota T, Kalyuzhniy O, McGuire A, et al. Rational HIV immunogen design to target specific germline B cell receptors. Science [Internet]. 2013 May 10;340(6133):711–6. Available from: http://www.ncbi.nlm.nih.gov/pubmed/23539181

63. Marcandalli J, Fiala B, Ols S, Perotti M, de van der Schueren W, Snijder J, et al. Induction of Potent Neutralizing Antibody Responses by a Designed Protein Nanoparticle Vaccine for Respiratory Syncytial Virus. Cell [Internet]. 2019;176(6):1420–1431.e17. Available from: http://www.ncbi.nlm.nih.gov/pubmed/30849373

64. Deng L, Mohan T, Chang TZ, Gonzalez GX, Wang Y, Kwon Y-M, et al. Double-layered protein nanoparticles induce broad protection against divergent influenza A viruses. Nat Commun [Internet]. 2018;9(1):359. Available from: http://www.ncbi.nlm.nih.gov/pubmed/29367723

65. Brouwer PJM, Brinkkemper M, Maisonnasse P, Dereuddre-Bosquet N, Grobben M, Claireaux M, et al. Two-component spike nanoparticle vaccine protects macaques from SARS-CoV-2 infection. Cell [Internet]. 2021;184(5):1188–1200.e19. Available from: http://www.ncbi.nlm.nih.gov/pubmed/33577765

66. Gu M, Torres JL, Li Y, Van Ry A, Greenhouse J, Wallace S, et al. One dose of COVID-19 nanoparticle vaccine REVC-128 protects against SARS-CoV-2 challenge at two weeks post-immunization. Emerg Microbes Infect [Internet]. 2021 Dec;10(1):2016–29. Available from: http://www.ncbi.nlm.nih.gov/pubmed/34651563

67. Keech C, Albert G, Cho I, Robertson A, Reed P, Neal S, et al. Phase 1-2 Trial of a SARS-CoV-2 Recombinant Spike Protein Nanoparticle Vaccine. N Engl J Med [Internet]. 2020;383(24):2320–32. Available from: http://www.ncbi.nlm.nih.gov/pubmed/32877576

68. Vu MN, Kelly HG, Kent SJ, Wheatley AK. Current and future nanoparticle vaccines for COVID-19. EBioMedicine [Internet]. 2021 Dec;74:103699. Available from: http://www.ncbi.nlm.nih.gov/pubmed/34801965

69. Malladi SK, Patel UR, Rajmani RS, Singh R, Pandey S, Kumar S, et al. Immunogenicity and Protective Efficacy of a Highly Thermotolerant, Trimeric SARS-CoV-2 Receptor Binding Domain Derivative. ACS Infect Dis [Internet]. 2021;7(8):2546–64. Available from: http://www.ncbi.nlm.nih.gov/pubmed/34260218

70. Ueda G, Antanasijevic A, Fallas JA, Sheffler W, Copps J, Ellis D, et al. Tailored design of protein nanoparticle scaffolds for multivalent presentation of viral glycoprotein antigens. Elife [Internet]. 2020;9. Available from: http://www.ncbi.nlm.nih.gov/pubmed/32748788

71. Antanasijevic A, Ueda G, Brouwer PJM, Copps J, Huang D, Allen JD, et al. Structural and functional evaluation of de novo-designed, two-component nanoparticle carriers for HIV Env trimer immunogens. PLoS Pathog [Internet]. 2020;16(8):e1008665. Available from: http://www.ncbi.nlm.nih.gov/pubmed/32780770

72. Chen X, Zaro JL, Shen W-C. Fusion protein linkers: property, design and functionality. Adv Drug Deliv Rev [Internet]. 2013 Oct;65(10):1357–69. Available from: http://www.ncbi.nlm.nih.gov/pubmed/23026637

73. van Rosmalen M, Krom M, Merkx M. Tuning the Flexibility of Glycine-Serine Linkers To Allow Rational Design of Multidomain Proteins. Biochemistry [Internet]. 2017 Dec 19;56(50):6565–74. Available from: http://www.ncbi.nlm.nih.gov/pubmed/29168376

74. Rawson S, Iadanza MG, Ranson NA, Muench SP. Methods to account for movement and flexibility in cryo-EM data processing. Methods [Internet]. 2016;100:35–41. Available from: http://www.ncbi.nlm.nih.gov/pubmed/27016144

75. Pang IK, Pillai PS, Iwasaki A. Efficient influenza A virus replication in the respiratory tract requires signals from TLR7 and RIG-I. Proc Natl Acad Sci U S A [Internet]. 2013 Aug 20;110(34):13910–5. Available from: http://www.ncbi.nlm.nih.gov/pubmed/23918369

