## Supplementary Figure S1-4 for "Comparative immunogenicity of bacterially expressed soluble trimers and nanoparticle displayed influenza hemagglutinin stem immunogens"

### Supplementary information

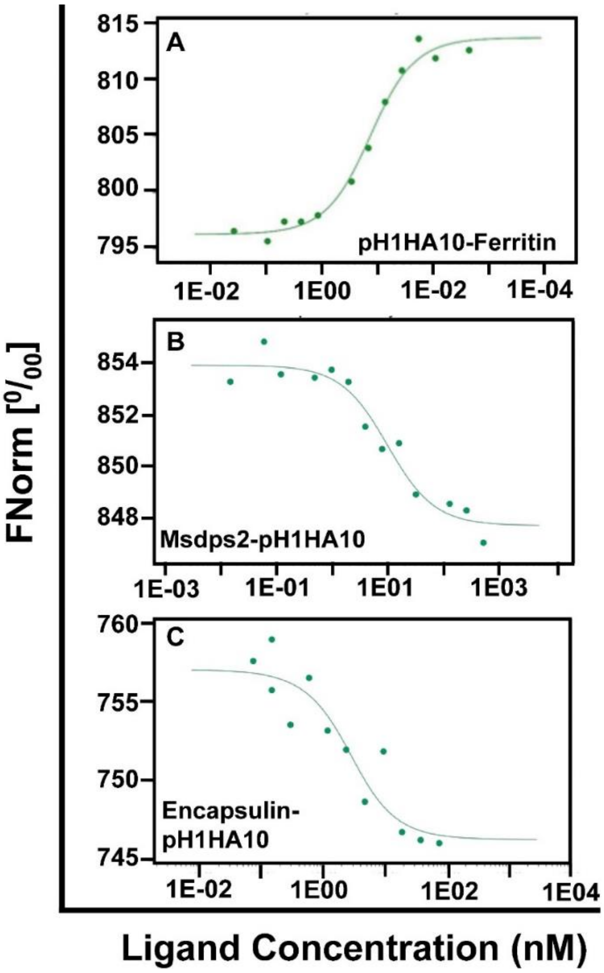


**Supplementary figure S1. MST of stem nanoparticles binding with bnAb CR6261**. CR6261 was labelled and used at a concentration of 100 nM. Non-fluorescent immunogens (A) MsDps2-pH1HA10, (B) Encapsulin-pH1HA10, and (C) pH1HA10-Ferrritin with concentrations ranging between 5 µM and 1 pM (in 1x PBS, pH: 7.4) were titrated and mixed with a fixed amount of labelled Ab, here CR6261. Normalized fluorescence FNorm [$^{0}/_{00}$] is plotted as a function of [ligand]. Dissociation constants ($K_D$) were analyzed using MO.Affinity Analysis software (version 2.2.5, NanoTemper Technologies) and are listed in Table 2.

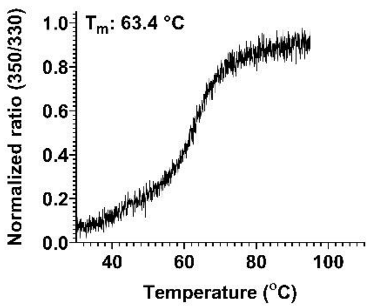


**Supplemental figure S2: Thermal denaturation profile of pH1HA10-Foldon using nanoDSF.** 2 µM of the protein in 1x PBS (pH 7.4) was incubated in the temperature range of 20 °C to 95 °C at 50% LED power.

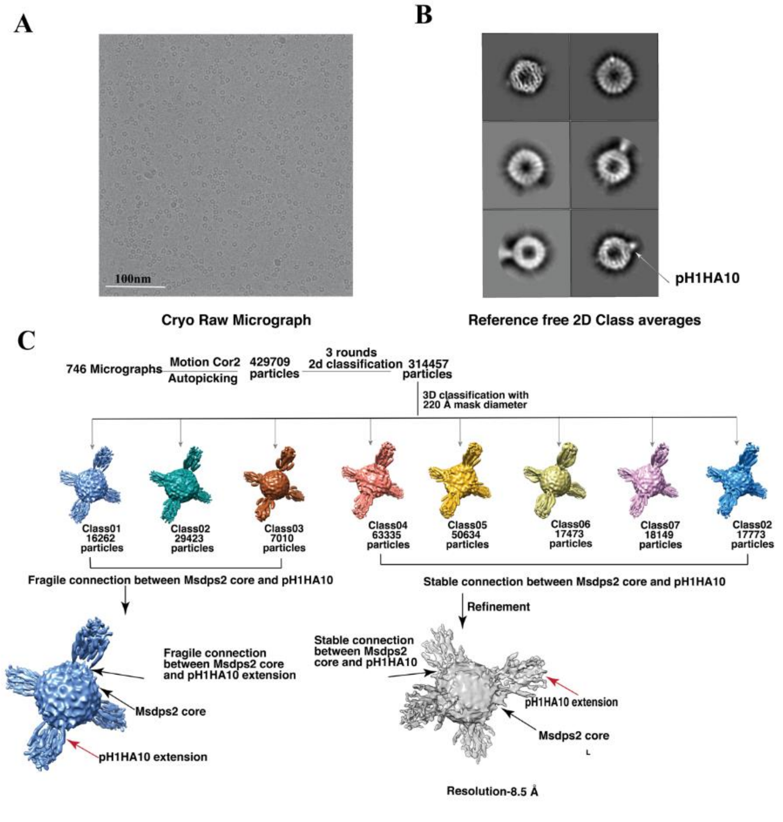


**Supplemental figure S3: 3D classification and structural characterization of Msdsps2-pH1HA10: (A)** Raw cryo-EM micrograph of Msdps2-pH1HA10**. (B**) Reference free 2D class averages of Msdps2-pH1HA10 nanoparticles**. (C)** Routine 3-D classification using 220Å particle diameter and 3D reconstruction of Msdps2-pH1HA10. A total of 746 micrographs were collected. Around 4,29,709 Msdps2-pH1HA10 particles were selected for structure determination. Three rounds of 2-D classification led to the identification of 3,14,457 distinct particles. The particles were then divided into eight classes. Classes having different connections (fragile or stable) between Msdps2 core and pH1HA10 extension were merged separately. The generated cryo-EM map obtained from the refinement of Msdps2- pH1HA10 particles with stable connections could be refined to a resolution of 8.5 Å.

A

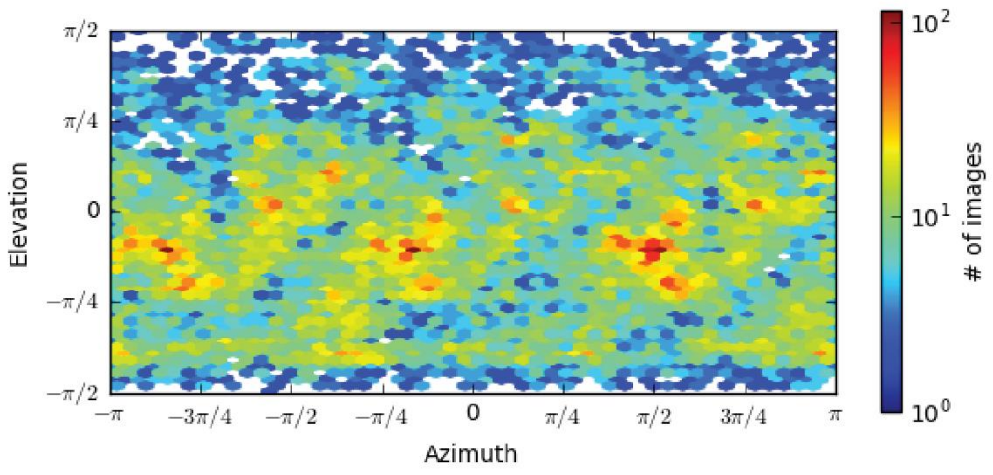


B

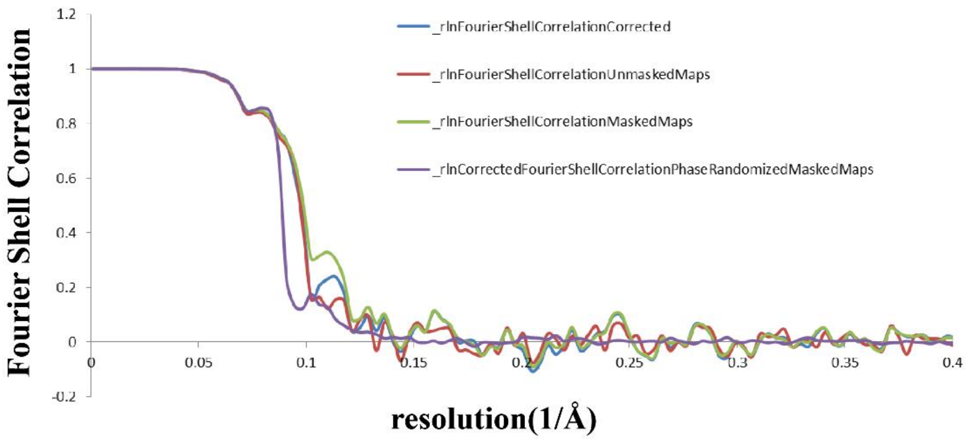


**Supplemental figure S4: (A)** Angular distribution of cryo-EM map of Msdps2-pH1HA10. **(**B) FSC curve of cryo-EM map of Msdps2-pH1HA10.
